# Inferring past effective population size from distributions of coalescent-times

**DOI:** 10.1101/033373

**Authors:** Lucie Gattepaille, Mattias Jakobsson

**Author notes:** Department of Evolutionary Biology, Norbyvagen 18D, 75 236 Uppsala, Sweden.

## Abstract

Inferring and understanding changes in effective population size over time is a major challenge for population genetics. Here we investigate some theoretical properties of random mating populations with varying size over time. In particular, we present an exact method to compute the population size as a function of time using the distributions of coalescent-times of samples of any size. This result reduces the problem of population size inference to a problem of estimating coalescent-time distributions. Using tree inference algorithms and genetic data, we can investigate the effects of a range of conditions associated with real data, for instance finite number of loci, sample size, mutation rate and presence of cryptic recombination. We show that our method requires at least a modest number of loci (10,000 or more) and that increasing the sample size from 2 to 10 greatly improves the inference whereas further increase in sample size only results in a modest improvement, even under as scenario of exponential growth. We also show that small amounts of recombination can lead to biased population size reconstruction when unaccounted for. The approach can handle large sample sizes and the computations are fast. We apply our method on human genomes from 4 populations and reconstruct population size profiles that are coherent with previous knowledge, including the Out-of-Africa bottleneck. Additionally, a potential difference in population size between African and non-African populations as early as 400 thousand years ago is uncovered.

---

Natural populations vary in size over time, sometimes drastically, like the bottleneck caused by the domestication of the dog (Lindblad-Toh *et al*. 2005) or the explosive growth of human populations in the past 2000 years (Cohen 1995). Inferring the demographic history of populations has various applications: it may lead to a better understanding of the impacts of major ecological or historical events (glacial periods (Lahr and Foley 2001; Palkopoulou *et al*. 2013), agricultural shifts or technological advances (Boserup 1981) and population admixture (Tishkoff *et al*. 2009; Schlebusch *et al*. 2012)). The demographic history should also be accounted for in studies of natural selection or in genome-wide association studies to avoid spurious results (Nielsen 2005; Marchini *et al*. 2004).

The particular problem of estimating past effective population size has gained considerable interest in recent years, in particular with the publication of methods such as the Bayesian skyline plots implemented in BEAST (Drummond *et al*. 2012) (see Ho and Shapiro (2011) for a review of this school of methods), and even more recently, methods such as PSMC (Li and Durbin 2011), MSMC (Schiffels and Durbin 2013) and DiCal (Sheehan *et al*. 2013). The former type of methods can use a rather large sample size, but can only account for a small number of loci. These methods have often been used solely for analyzing mitochondrial DNA. The latter group of methods are designed to handle genome-wide data and explicitly model recombination using a Markovian assumption for neighboring gene-genealogies (McVean and Cardin 2005). PSMC works with a single (diploid) individual, which lead to simple underlying tree topologies without requiring phase information. However, the inference power in the recent past is limited, as most coalescences in a sample of size 2 are not expected to occur in the recent past (Li and Durbin 2011). MSMC and DiCal extend this approach by using information from multiple samples. MSMC focuses on the first coalescence event in the sample at each locus and ignores the remaining coalescence events. The algorithm can deal with genome-wide data in a computationally efficient way. DiCal, on the other hand, uses all coalescent events in the gene-genealogies to provide estimates of the population size, assuming a Markov property between sites as well (Sheehan *et al*. 2013). The algorithm quickly becomes computationally intensive as the sample size increases, thus a genome-wide use of the method is currently still limited.

There are two important steps for most of these types of approaches: the inference of the underlying gene-genealogies, and the inference of population size as a function of time from the inferred genealogies. In this paper we introduce the **Pop**ulation **Si**ze **C**oa**l**escent-times based **E**stimator (Popsicle), an analytical method for solving the second part of the problem. We derive the relationship between the population size as a function of time and the coalescent-time-distributions by inverting the relationship of the coalescent-time-distributions and population size derived by Polanski and colleagues (Polanski *et al*. 2003), where they expressed the distribution of coalescent-times as linear combinations of a family of functions that we describe below. Using a simple algorithm of tree inference from genetic data, we can investigate the effects of a range of conditions associated with real data such as finite number of loci, sample size, frequency of mutation and presence of cryptic recombination. We apply Popsicle to sequence data from the 1000 genomes Project and uncover population size histories that are coherent with general knowledge of human past effective population size.

## From the distributions of coalescent-times to the population size

Under the constant population size model, the waiting times *T_n_*, *T_n_*_−1_, ⋯, *T*_2_ between coalescent events are independent exponentially distributed random variables. In particular, the time *T_k_* during which there are exactly *k* lineages follows an exponential distribution with rate 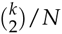 generations. When population size varies as a function of time (*N* = *N*(*t*)), the waiting times to coalescence are not independent any longer. Specifically, for *k* ∈ [2, *n* − 1], *T_k_* depends on all the previous coalescent-times from *T_k_*_+1_ to *T_n_* (see *e.g.* Wakeley (2009) for an extensive description of the coalescent).

In this paper, we derive a relationship between *N*(*t*) and the distributions of the *cumulative* coalescent-times, that we denote by *V*. More specifically, for *k* ∈ [2, *n*]:

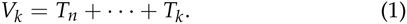

The *V_k_* variables represent the sum of times from present to each coalescent event. Because we only use the cumulative coalescent-times *V_k_* and not the individual times *T_k_,* we refer to as *coalescent-times* the times *V_k_* for *k* ∈ [2, *n*], omitting the term *cumulative* for convenience. For example, the random variable *V*_2_ represents the time to the Most Recent Common Ancestor (T_MRCA_). All coalescent-times *V_k_* are expressed in generations. We denote as *π_k_* the density function of *V_k_*.

Polanski and colleagues (Polanski *et al*. 2003) derived the density function of coalescent-times under varying population size as linear combinations of a set of functions 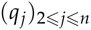, where

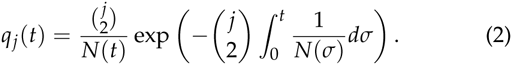

Similar functions have previously been used in a context of varying population size (Griffiths and Tavare 1994). For *k* ∈ [2, *n*], the relationship between the density function *π_k_* and 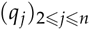

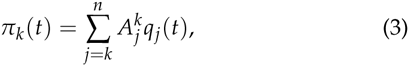

with

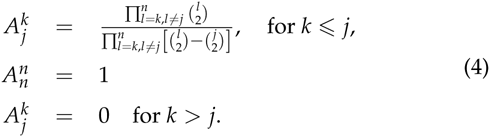

We also define the integral of *q_j_* with respect to t as

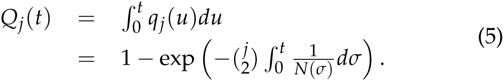

From equations 2 and 5 we can derive that:

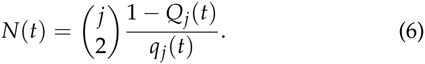

The principle of our method is to use the distributions of the coalescent-times to get to the *q_j_* functions. In other words, we invert the result of Polanski *et al*. (2003).

### Theorem 1.

Given a sample of size *n*,

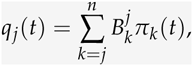

with

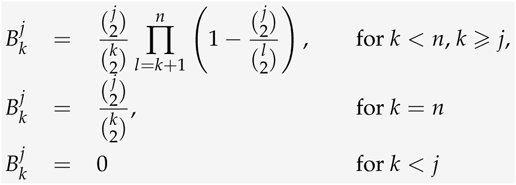

The proof of the theorem is given in the Supporting Information. This theorem implies that for any time *t* generations in the past, *q_j_*(*t*) and *Q_j_*(*t*) can be obtained using the distributions of coalescent-times. From each *q_j_* (and its integral *Q_j_*), the function *N*(*t*) can be obtained using equation 6. In contrast to the 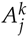 coefficients (equation 4) which can become very large as *n* increases and are of alternate signs (Polanski *et al*. 2003), the 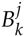 coefficients introduced in theorem 1 are stable, positive and take values between 0 and 1 (Fig. 1). Thus, our formula is not constrained by numerical limitations and can be used for very large sample sizes.

**Figure 1.**
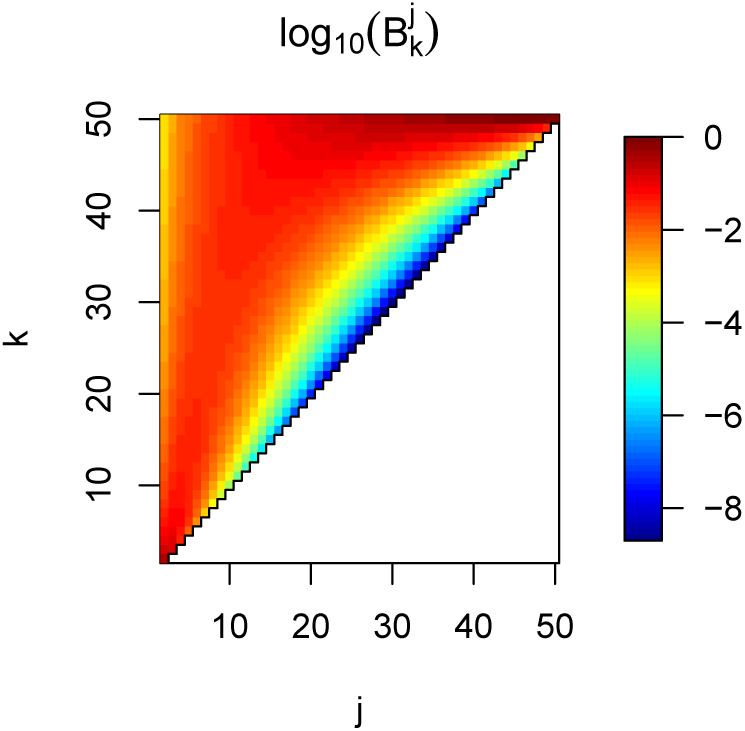
**Heatmap of the values of** 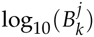 **for** *n* = 50**, as function of** *k* **and** *j*. The white area represents the region where 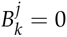.

## Application of the theorem on simulated gene-genealogies

Theorem 1 states that the population size can be computed at any time in the past, provided that we know all the *n* − 1 distributions of coalescent-times for any time in the past. However, this knowledge would require to observe the genealogies of an infinite number of independent loci evolving under the same *N* function over time. In practice, genomes are finite so we only have access to a finite number of loci to estimate of the coalescent-time distributions. We use empirical distribution functions 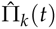 to estimate the cumulative distribution functions Π*_k_*(*t*) of the coalescent-times as these estimators have good statistical properties: they are unbiased and asymptotically consistent Van der Vaart (2000). From Theorem 1, we estimate the *Q_j_* functions by

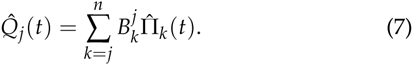

Because of the finite number of loci, time is discretized into intervals and *N*(*t*) within each interval is estimated by its harmonic mean, as the harmonic mean of *N* has a simple relationship to the *Q_j_* functions:

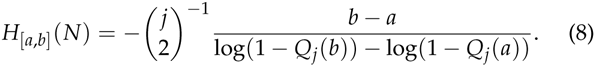

In the remainder of the paper, we set *j* = 2 (in equations 7 and 8), as it incorporates information from all coalescent-time distributions and perform well even for very recent times (see Supporting Information Figures S1, S2 and S3).

### Evaluation of N(t) inference

In order to evaluate the inference of *N*(*t*), we used the software ms (Hudson 2002) to simulate samples under different population models with varying population size. We investigated 4 demographic scenarii illustrated in Fig. 2. The first 3 scenarii describe demographic models that span between the present and 100,000 generations in the past and that include various periods of constant population size, instantaneous changes, exponential growth or decline. In contrast to scenarii 1, 2 and 3, scenario 4 describes changes in size that occur in the recent past, within the last 2,000 generations. Detailed descriptions of each scenario and the ms commands for the simulations are given in the Supporting Information (Section 2 and tables S1, S2, S3 and S4). In each studied scenario, we simulated 1,000,000 independent gene-genealogies of 20 haploid gene-copies (note that we will investigate the effect of number of loci and hence reduce that number for certain cases, see below). We assume that the true gene-genealogies are known and omit any inference of genealogies from polymorphism data at this stage. The genealogies were used to estimate coalescent-time distributions and in turn reconstruct the population size profile using theorem 1. We discretized time into 100 equally long intervals (1,000 generations in each interval for scenarii 1, 2 and 3, and 20 generations in each interval for scenario 4).

**Figure 2.**
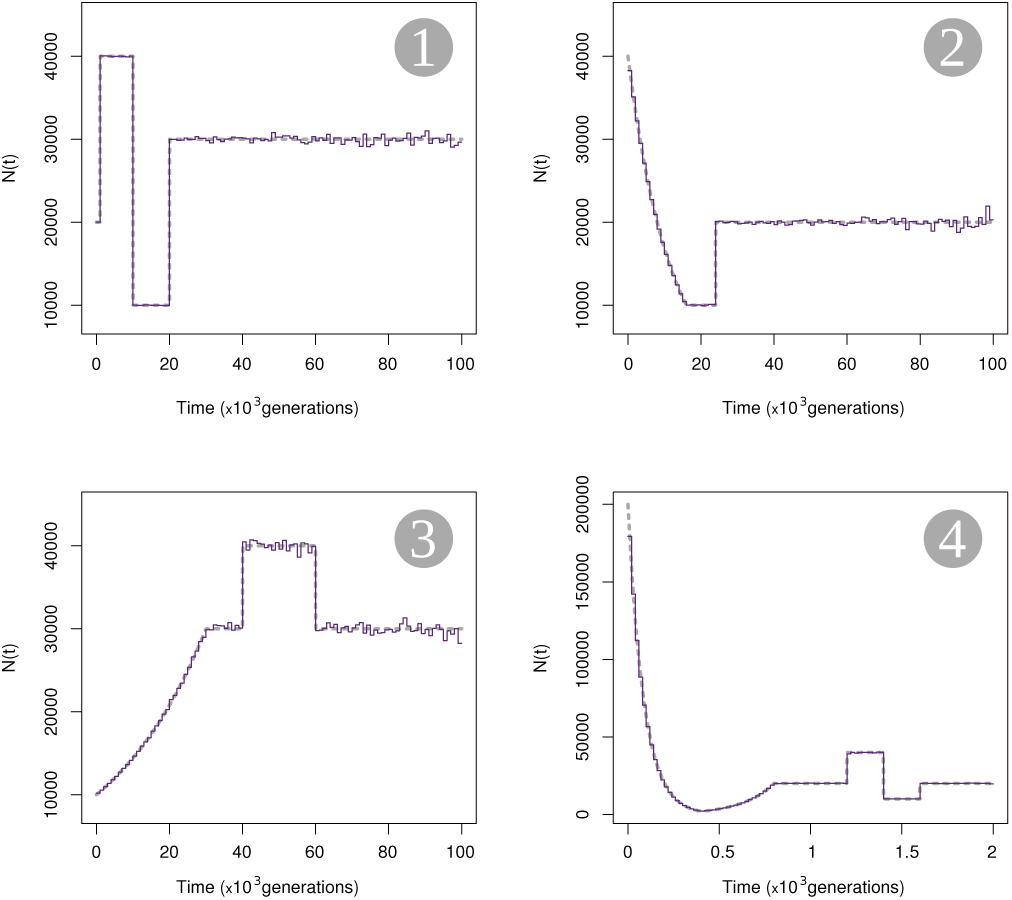
Estimation of *N* based on simulated gene-genealogies. Four scenarii of variable population sizes are used to generate 1,000,000 independent loci in each scenario, for a sample of size 20 (10 diploid individuals). Time is divided into 100 regular intervals and estimates of the harmonic mean of *N* (purple solid lines) for all intervals are plotted. The true values of *N* over time are indicated by gray dashed lines.

The harmonic mean estimates are very close to the true size in all 4 scenarii, with better accuracy in the recent past than in the distant past (Fig. 2). The division of time into 100 intervals is arbitrary and dividing time using the true break points of the scenarii leads to an almost perfect fit for the time periods where the population size is constant, whereas dividing time more finely in the periods of variable size improves the estimation, as long as there are enough coalescent-times occurring within the interval to get a good estimate of the cumulative distribution function (results not shown). The *N*(*t*) estimation is very accurate in periods of small population size, especially when it is followed by an expansion. Estimates of *N*(*t*) are more variable around the true value when population size was larger in the past (scenario 3). These observations can be understood intuitively by the fact that *π*(*t*) will be better estimated in time-periods of small *N* as the coalescence rate is proportional to the inverse of N. The resolution of the reconstruction method for *N* is also accurate in the recent past, even for drastic or rapid changes in size over a couple of hundred generations (scenario 4). In summary, with a finite but sufficiently large number of loci to estimate the cumulative distributions of coalescent-times, we can accurately reconstruct the global shape of the population size over time, from very recent times to far into the past.

### Effect of sample size

We tested the accuracy of our method for different sample sizes. To be able to quantify the performance in reconstructing the population size over time, we introduced two statistics: the *average relative difference* (ARD), and the *average relative error* (ARE). The former quantifies a systematic deviation from the true value of the population size, while the latter quantifies the error of the estimation (see Methods section for the computation of ARD and ARE). We used scenarii 1 and 4, from which we simulated 1,000,000 independent gene-genealogies with sample sizes taken from the values {2, 5, 10, 15, 20, 30}. Each scenario was divided into large periods, to be able to discriminate the effect of sample size in the *N*(*t*) reconstruction between recent and old time periods and between periods of large and small population sizes. Scenario 1 was divided into 5 periods, while scenario 4 was divided into 6 periods (table 1, Fig. 3). Within each period, we discretized the time into 100 equally long intervals and assessed the *N*(*t*) reconstruction with ARD and ARE (Fig. 3).

**Table 1.** Division of scenarii 1 and 4. Time intervals are given in generations.

| Period | Scenario |  |
| --- | --- | --- |
|  | 1 | 4 |
| 1 | [0-1,000] | [0-400] |
| 2 | [1,000-10,000] | [400-800] |
| 3 | [10,000-20,000] | [800-1,200] |
| 4 | [20,000-60,000] | [1,200-1,400] |
| 5 | [60,000-100,000] | [1,400-1,600] |
| 6 | - | [1,600-2,000] |

In general, scenario 1 is predicted more accurately than scenario 4, with an average relative error ranging between 0.2% and 3.6% compared to a range of 0.4% to 12.1% for scenario 4. There is little bias in the reconstruction of the 2 scenarii, except maybe for sample of size 2 in scenario 4, where there may be an upward bias of some 5% in period 4. In both scenarii and in all periods, the accuracy of the estimates are improved by increasing the sample size. The improvement is substantial when increasing the sample size from 2 to 10 and increasing the sample size further only results in modest improvements. Note the relatively higher error for the instantaneous population expansion of scenario 4 (period 4), irrespective of sample size, suggesting that a large population size for a brief period of time is difficult to infer. Accurate estimates of *N* for such periods require a greater number of loci to obtain resolution on par with time-periods with smaller *N*, as the number of coalescences is reduced for periods of large *N*. This effect is investigated further in the next section.

**Figure 3.**
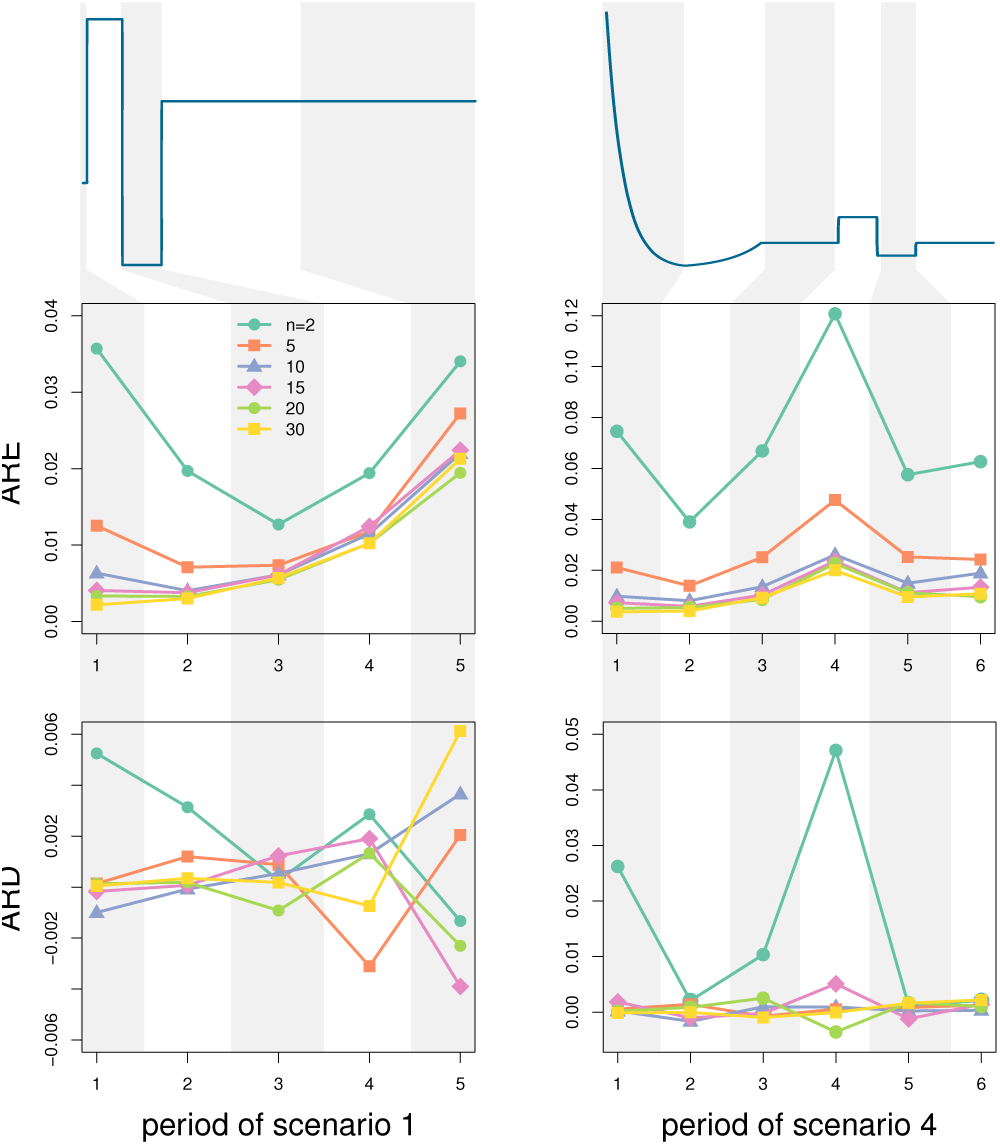
Effect of sample size. We divide scenario 1 (left panel) and scenario 4 (right panel) into smaller periods of time were we assess the average relative error and the average relative difference on the *N* estimates compared to the true values of *N*, as functions of the sample size used for the estimation. 1,000,000 loci were simulated for each scenario and each sample size. The original scaling for the x and y axis of both scenarii can be found in Fig. 2.

### Effect of the number of loci

With the full knowledge of the density functions *π_k_*, we could potentially compute *N* at any time in the past. However, in practice, the distributions can only be estimated where observations are made, hence we are limited to the time ranges where reasonable estimates of the distributions can be computed because we have enough observations. For that reason, the more loci, the more coalescent-times can be observed within a time interval and the better the estimate of the cumulative distributions. Here we investigate the robustness of Popsicle to varying number of loci, by simulating genealogies of samples of size 20 under scenarii 1 and 4. We compare the effect of the number of loci for different periods in the past, as described in table 1, divide each period into 100 regular intervals on which we estimate the harmonic mean of *N* and measure the accuracy within each period with ARD and ARE.

As expected, the accuracy of the *N* estimates in all periods for both scenarii increases with increasing number of loci (fig. 4). For small numbers of loci, errors can reach 40% and more. For scenario 1 and 1,000 loci, no coalescence occurred during periods 4 and 5 in any of the simulations, making the inference impossible for these periods. Similarly, there were no coalescence events in period 5 of scenario 1 with 5,000 loci, as well as periods 4 and 6 in scenario 4 with 1,000 loci. This illustrates the greater difficulty of accurate *N* reconstruction for older periods of time, and periods of large population size, both subject to low probabilities of observing coalescences. Thus, depending on the history of the population and how far back in time *N* is of interest, the required number of loci will vary. Sub-sampling from some particular number of loci might give an idea on whether or not that number is enough for a good estimation of *N* over time.

**Figure 4.**
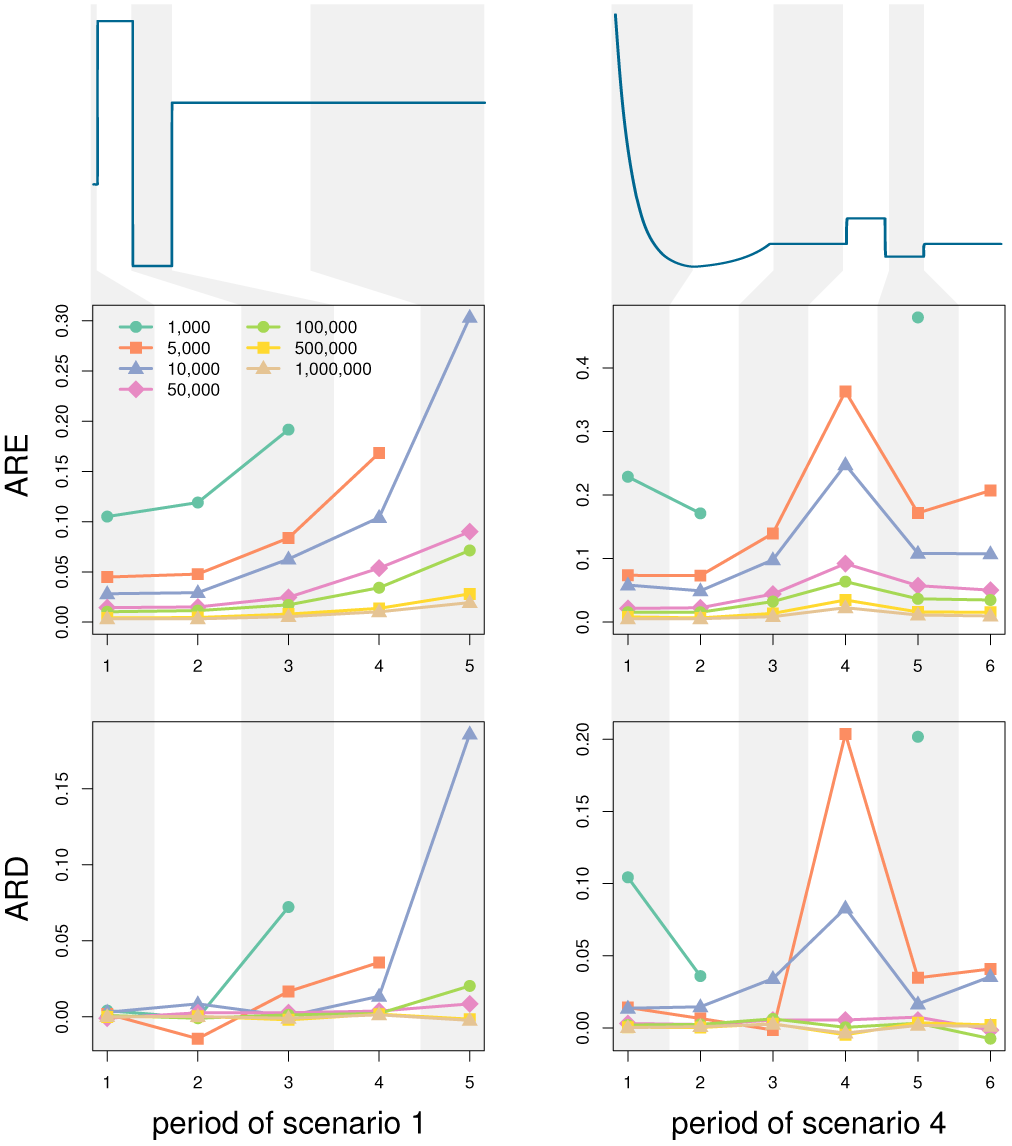
Effect of the number of loci. We divide scenario 1 (left panel) and scenario 4 (right panel) into smaller periods of time were we assess the average relative error and the average relative difference on the *N* estimates compared to the true values of *N*, as functions of the number of loci used for the estimation. The sample size for all simulations is 20. The original scaling for the x and y axis of both scenarii can be found in figure 2.

## Discussion

In this paper, we derived the analytical relationships existing between the population size over time of a randomly mating population and the distributions of coalescent-times. We showed by simulations that with a sufficient number of loci and their underlying gene-genealogies, we can reconstruct *N*(*t*) accurately at various time scales, with a greater accuracy in the recent past and in periods of smaller population size (Fig. 4, Supporting Information Figure S4). The results also showed an improvement in the reconstruction of *N*(*t*) with an increase in sample size, however the improvement becomes modest for sample sizes larger than 15 haploid gene-copies.

The major implication of our main result is to reduce the problem of *N*(*t*) reconstruction from polymorphism data to a problem of gene-genealogy inference. If local gene-genealogies in the genome can be inferred accurately from observed polymorphism data, then our theorem can be used to estimate *N*(*t*) with great accuracy as well. Currently, however, local gene-genealogy inference remains a challenge. First, most genomes do not consist in large sets of independent non-recombining loci, but rather in sets of recombining chromosomes. Each chromosome can be seen as a linear structure of successive non-recombining loci whose underlying genealogies are correlated with one another. This correlation decays with distance between loci due to recombination. Also, in a given sample, the exact positions on the chromosome of the recombination events, hence the break points between the non-recombining bits of DNA, are unknown. Fully recovering the genealogies along the chromosome means reconstructing the ancestral recombination graph from polymorphism data and this is a challenging problem that has drawn much attention in the last decades (Griffiths and Marjoram 1996; McVean and Cardin 2005; Parida *et al*. 2008; Zheng *et al*. 2014; Rasmussen *et al*. 2014). Ignoring recombination and treating recombining loci as non-recombining can lead to the inference of spurious changes in population size, even under the simple model of constant population size (Fig. 5, Supporting Information Figure S5), because of the effect it has on genealogies. In particular, genealogies inferred from recombining loci are weighted averages of the underlying genealogies of the non-recombining fragments of the loci, and therefore tend to be more star-like as well as of intermediate size. Estimating one single gene-genealogy from such a mosaic of correlated gene-genealogies will have an impact on the distributions of coalescent-times (see Supporting Information Figure S6).

For some species, there might be simply not enough mutation events to be able to infer the local gene-genealogies of non-recombining segments. In humans for example, the ratio between the mutation rate per site and per generation and the recombination rate per site and per generation is likely close to 1 (or 2, depending on the mutation rate assumed, the pedigree based mutation rate or the divergence based mutation rate, see *e.g.* Scally and Durbin (2012)). Hence, on average, for each mutation observed locally in a sample, there is also a recombination break-point nearby. We assessed the effect of the mutation rate per locus by using a simple algorithm of gene-genealogy inference from polymorphism data and applied Popsicle on the obtained distributions of coalescent-times (Fig. 6, see also Methods section for a description of the employed algorithm). The mutation rate is important for an accurate reconstruction of *N*(*t*): the higher the mutation rate, the more information there is to infer the gene-genealogies and the better is the reconstruction of *N*(*t*). This effect is particularly important for recent times where enough mutations are required to accumulate to infer the very recent population sizes (S7 Fig).

### Application to the 1000 Genomes sequence data

Despite the challenges of inferring gene-genealogies discussed above, it is possible to apply our method on empirical sequence data. The effect of recombination can be mitigated by considering only regions of the genome with low or no recombination, provided that we have access to a good genetic map. Following this principle, we applied Popsicle to human genome sequence data from the 1000 Genomes Project (Complete Genomics high coverage samples from the (Complete Genomics data from 1000 Genomes public repository 2013)), for Yoruba individuals from Nigeria, for American individuals of European ancestry from Utah, U.S.A, for Han Chinese individuals from southern China and for Peruvian individuals. We extracted regions of no recombination according to the Decode recombination map (Kong *et al*. 2002) (see Methods section for a description of the data preparation). We inferred *N*(*t*) profiles for the 4 populations in two ways: a) using single individuals (as PSMC does) and averaging across individuals (denoted ‘Popsicle 1’), and b) using 5 individuals at once from the population and averaging across the the samples of individuals (denoted ‘Popsicle 5’).

**Figure 5.**
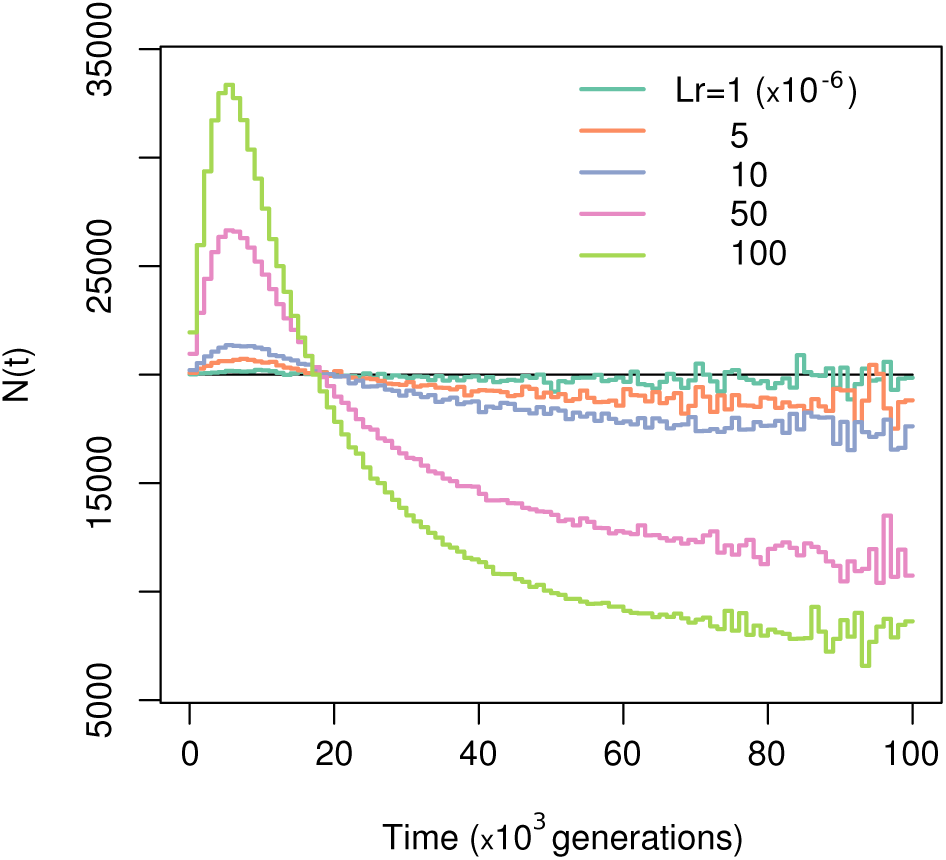
Effect of ignoring recombination. Comparison between *N*(*t*) reconstructed using gene-genealogies computed as a weighted average of the gene-genealogies obtained from ms and true *N*(*t*) (black lines) under a model of constant population size. We generated 1,000,000 independent loci for 20 haploid gene-copies for 5 different levels of recombination within each locus. The different cryptic recombination rates for each locus (in Morgan) is indicated by different colors and the values of the recombination of the segments are given in the legend. Assuming a recombination rate *r* per site and per generation of 1.25 × 10^−8^, the considered sequences are 80, 400, 800, 4000 and 8000 base pairs long.

Overall, the Popsicle profiles of effective population size in the last million years for every population largely resemble the vague knowledge about past human population sizes as well as the *N*(*t*) profiles inferred by *e.g.* PSMC (Fig. 7A). For instance, Popsicle reveals a steady but slow increase in effective population size staring around 1 million years ago, reaching a maximum between 200,000 and 500,000 years ago, followed by a sharper decline and a recovery during the last 100,000 years for European and East Asian populations. However, prior to a million years ago, the population size inferred by PSMC is higher than the population size inferred by Popsicle (Supporting Information Figure S8). In addition, Popsicle infers a less sharp decline in population size than PSMC does, for all four populations, and infers a population size history markedly different for Yoruba compared to the three other non-African populations (Fig. 7B and Fig. 7C) whereas the Yoruban population follows the non-African populations rather closely in the PSMC results (Supporting Information Figure S9). Popsicle results suggest a somewhat larger ancestral population for Yoruba than the ancestral population size of the 3 non-African populations, which could be interpreted as deep and long-lasting population structure within Africa between 400,000 and 100,000 years ago. Note however that the non-recombining regions have been chosen using the Decode recombination map, a genetic map formed by tracking more than 2,000 meioses in Islandic lineages. Recombination patterns and hotspots in particular are believed to be variable across populations (Myers *et al*. 2005; Baudat *et al*. 2010), thus the non-recombining regions selected using the Decode map might be in fact recombining in Yoruba, resulting in a bias of the population size estimates (see Fig. 5). Recombination maps for Yoruba have been computed (Frazer *et al*. 2007), but because they have been inferred using properties of linkage disequilibrium which itself depends on demography, they are cannot be used in this context. A future pedigree- or sperm-typing-based recombination map for the Yoruba would likely resolve the differently inferred *N*(*t*) profiles for African and non-African populations.

**Figure 6.**
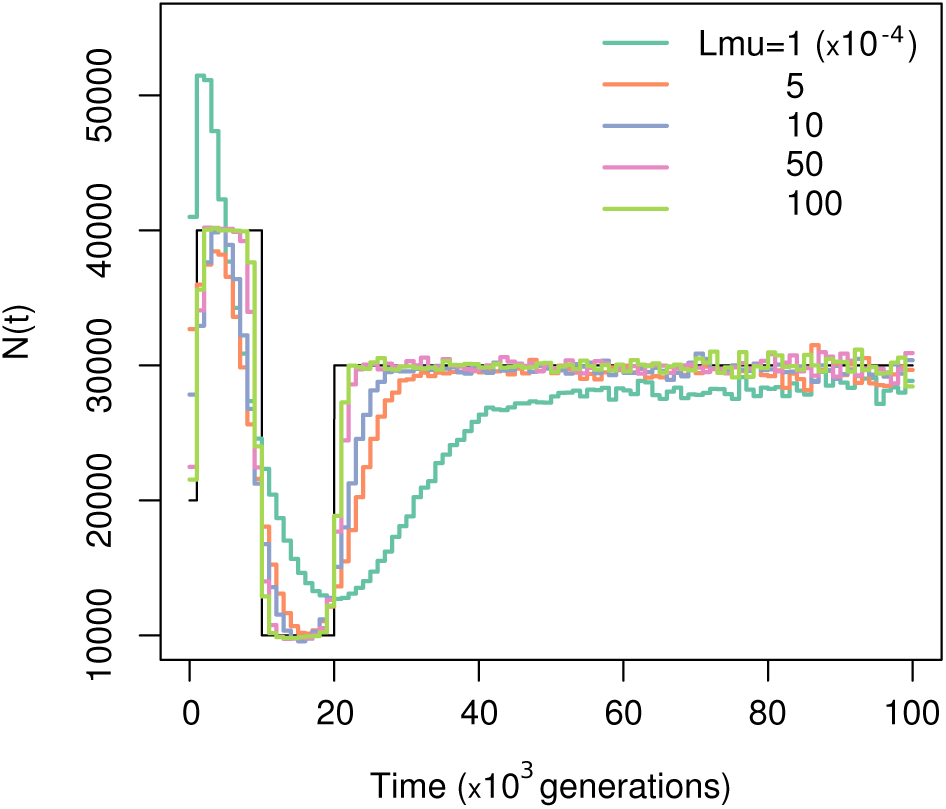
Effect of estimating gene-genealogies from polymorphism data. Reconstruction of *N*(*t*) from distributions of coalescent-times computed from gene-genealogies inferred from polymorphism data. We used a sample size of 20 and 1,000,000 independent loci, evolving under scenario 1. The mutation rate per locus *Lµ* is indicated by the color of the line and the legend gives the mutation rates.

Popsicle 1 and Popsicle 5 give similar effective population size profiles (Fig. 7B and 7C) but the time of the major features in Popsicle 5 are shifted to older times compared to Popsicle 1. Whereas Popsicle 1 suggests a bottleneck in non-African populations that reaches its strongest effect between 30,000 and 40,000 years ago, Popsicle 5 places the bottleneck between 70,000 and 80,000 years ago, which is more in line with the estimates of timing of the founder effects due to a dispersal out-of-Africa (Scally and Durbin 2012). In none of the applied methods (Popsicle or PSMC) do we see the super-exponential increase in size that has occurred in all populations since the spread of agriculture (Keinan and Clark 2012). It is possible that too few loci are included for a reliable inference in the recent times or that the mutation rate per locus does not allow us the required time resolution to observe this phenomenon, as very recent gene-genealogies are likely to contain no mutation thus no information if the mutation rate is too low. Another possible explanation could be that we cannot observe enough rare variants with only 22,321 loci, which has been recently deemed necessary to observe the exponential growth human populations have been going through in the past thousands of years (Keinan and Clark 2012).

**Figure 7.**
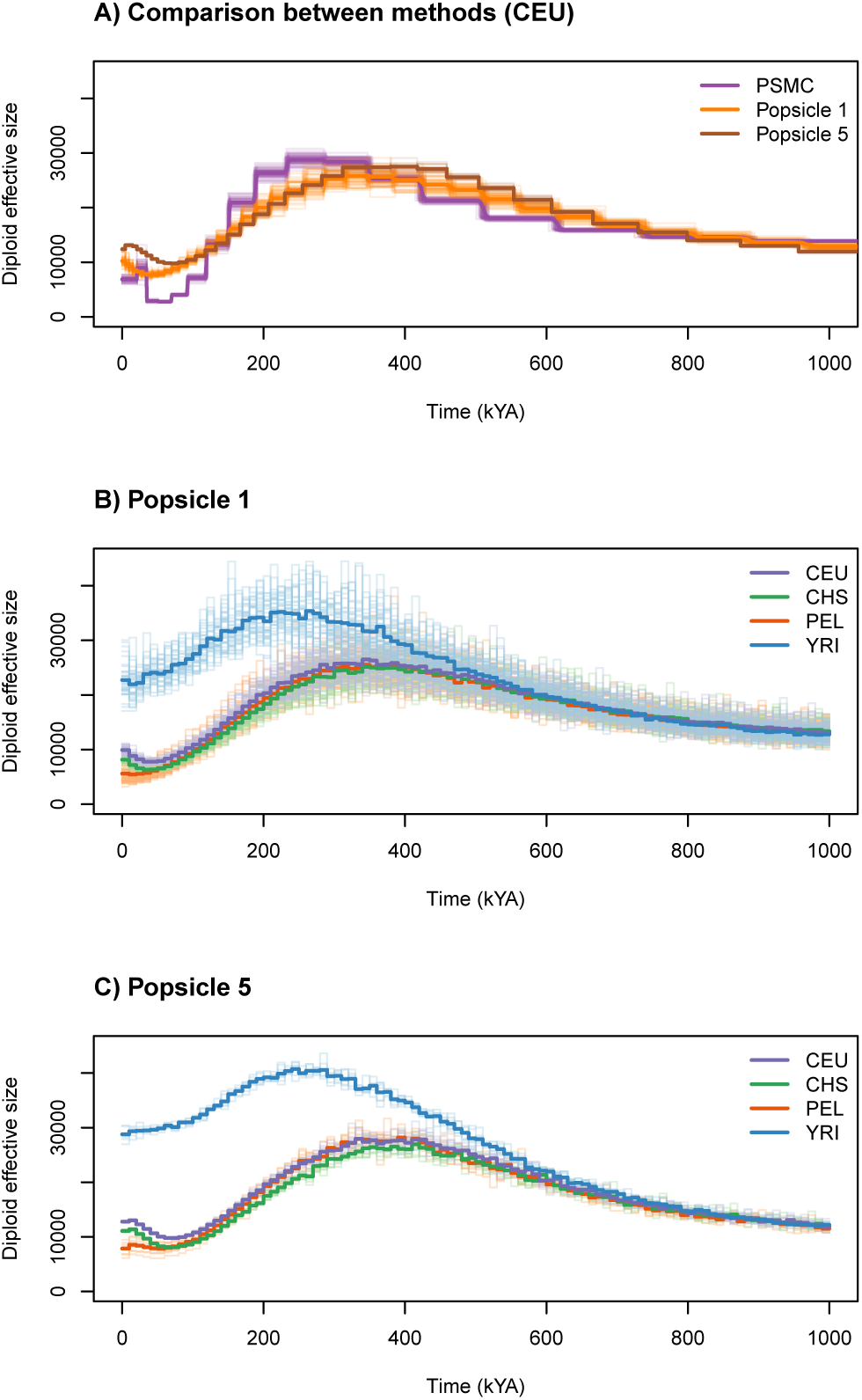
Comparison of *N*(*t*) between CEU, CHS, PEL and YRI. The time scale is computed assuming a mutation rate of 1.25 × 10^−8^ and a generation time of 25 years.

The resolution of Popsicle can be better than that of PSMC, as Popsicle does not constrain the coalescent-times into a finite (and usually rather small) set of values like PSMC does. In principle, any time discretization for computing the harmonic mean of the effective population size over time can be used, though in practice we need to make sure that there are enough coalescences within each time interval to get reliable estimates of the effective size. Popsicle is also markedly faster than PSMC, not only because it uses a low number of non-recombining regions, but also because of the closed form relationship between population size and coalescent-time distributions. Most of the computational time is spent on inferring the gene-genealogies (which takes less than 20 minutes for the 22,321 loci in the data application). Once the gene-genealogies are computed, the application of the theorem for reconstructing the population size takes a few seconds. Finally, Popsicle accommodates for samples of any size, which should lead to more reliable results, especially in the recent times, provided that the phasing of the genomes is accurate.

Applying Popsicle on extracted regions of limited recombination should not bias the results in principle. Regardless of the molecular reason explaining the low rate of recombination in the region (for instance, limited access for crossovers or conservation constraints due to functional importance of the region), the fact that there is one local gene-genealogy for the entire region is what matters for the method to work. However for applications to empirical data, variation in the local mutation rate, due to purifying selection for example, will affect the reconstruction of the gene-genealogy by changing the estimates of the branch lengths for different loci. This could potentially cause bias in the reconstructed Popsicle profiles, as all gene-genealogies are inferred using one mutation rate. Using a mutation map obtained from the study of *de novo* mutations in trios or pedigrees could alleviate this issue and infer the local gene-genealogies from genetic data using a specific mutation rate for each region.

### Conclusion

We present a novel method for inferring population size over time, a problem that has recently gained great attention due to the availability of genome data. By analytically relating *N*(*t*) to the distribution of coalescent-times, we have connected *N*(*t*) to the problem of inferring the ancestral recombination graph from polymorphism data, which remains a challenge in population genetics. We showed that, even using a moderate number of loci and a simple algorithm for genealogy inference, our method Popsicle was able to recover the general pattern of population size as a function of time, which have been observed previously using other methods, but with greater resolution and faster computational time, which will be useful for future large-scale genome studies.

## Methods

### Quantifying the accuracy of the method

Let’s consider a time discretization (*t*_0_, *t*_1_, ⋯, *t_m_*), then we define ARD and ARE are defined as:

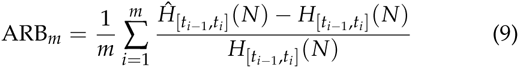

and

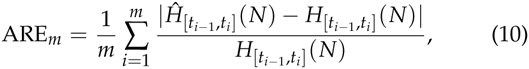

where 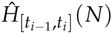 is the estimate of the harmonic mean of *N* during the time interval [*t_i_*_−1_, *t_i_*] as defined in equation 8 with *j* = 2 and *Q*_2_ replaced by its estimate 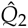, whereas 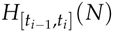 is the value for the true harmonic mean of *N* on the corresponding interval.

### Effect of ignoring recombination

We explore the robustness of the *N*(*t*) reconstruction when recombination occurs in the loci but each locus is treated as non-recombining. Instead of considering the multiplicity of trees within one locus, we consider the entire segment as non-recombining and having a single genealogy, represented by an average tree. The true nature of this “average tree” can be difficult to define when the sample size is strictly greater than 2, however for a sample of size 2, the average tree is simply the weighted mean of the trees of the non-recombining segments, with the weight being the relative length of each non-recombining segment compared to the total segment length. We investigate the effect of ignoring recombination for samples of size 2 and for samples of size 20, for different levels of recombination within each simulated locus. For the samples of size 20, we build the “average tree” by applying a UPGMA algorithm on the weighted average matrix of pairwise time to coalescence between all pairs of haploid individuals. We use scenarii 1 and 4, as well as the constant size model, to study the robustness of the method to different levels of recombination. We tested 5 levels of recombination within the locus: 10^−6^, 5 × 10^−6^, 10^−5^, 5 × 10^−5^ and 10^−4^. Assuming a recombination rate of 1.25 × 10^−8^ per site per generation, which is around the estimated average of the human recombination rate, these 5 levels represent loci of length 80, 400, 800, 4,000 and 8,000 base pairs.

### Inferring gene-genealogies from polymorphism data

We apply a simple 2-step algorithm to infer gene-genealogies from polymorphism data. In the first step, we naively reconstruct the genealogy for each locus using the UPGMA algorithm on the matrix of pairwise differences then convert the branch lengths from a time scale in mutations to a time scale in generations using the mutation rate per locus, which is considered known. Because of the discrete behavior of mutation, we do not really have resolution at this stage for time intervals smaller than 1/(2*Lµ*) generations, with *Lµ* being the total mutation rate of each locus. So, we discretize the time space into equal intervals of size 1/(2*Lµ*), starting at 0 and estimate the harmonic mean of *N* on each interval using the method. This gives us a first view on the profile of the population size over time. In the second step, we refine our reconstruction by using the *N*(*t*) profile computed in the first step. More precisely, we now use the pairwise differences between haploid individuals to estimate the time to the most recent common ancestor of each pair of individuals and thus define a distance matrix on which we apply UPGMA to reconstruct the genealogy. We compute the coalescent-times between the pairs using a Gamma distribution, following the idea that if mutations are Poisson distributed onto the coalescent tree of a given pair, and if the height of the tree is exponentially distributed with rate 1/*N_e_* (which is the case under the constant model of *N*(*t*)= *N_e_*), then the height of the tree *T*, conditional on the number of pairwise differences *S* between the two individuals, is Gamma distributed with shape *S* + 1 and with rate 2*Lµ* + 1/*N_e_* Tavaré *et al*. (1997):

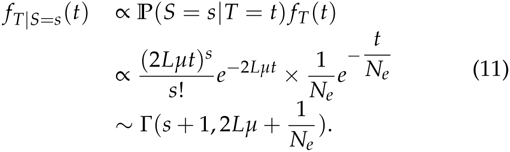

We use the first step to compute *N_e_* as the harmonic mean of the inferred *N* from present to the time interval corresponding to the number of observed differences between the two individuals. We present the results of the described protocol applied on samples of size 20, simulated with values of *Lµ* taken from {10^−4^, 5 × 10^−4^, 10^−3^, 5 × 10^−3^, 10^−2^}, for 1,000,000 loci and under scenarii 1 and 4 (Fig. 6 and S7 Fig). For reference, with a mutation rate of 1.25 × 10^−8^ per base pair per generation, the range of *Lµ* values corresponds to loci of 8, 40, 80, 400 and 800 kb respectively.

### Application to Human data

**Data preparation.** We use high coverage sequencing data from the 1000 Genomes Project, publicly available at ftp://ftp.1000genomes.ebi.ac.uk/vol1/ftp/data/. The data is downloaded as VCF formatted files, from which we retain variant positions passing the filters set up by the 1000 Genomes Project, replacing the filtered out positions by missing genotypes. We retain only the trios and within each population existing in the sample, we phased the individuals using BEAGLE Browning and Browning (2007) under the trios file input option, but retain only the parents after phasing, as a sample of unrelated individuals. In doing the phasing, BEAGLE also imputes missing genotypes. We extract sequences corresponding to regions of supposedly no recombination as indicated by a recombination rate of 0 in the Decode genetic map Kong *et al*. (2002). The description of how those regions were ascertained is given below. The populations we use in our study are CEU (individuals of European ancestry from Utah, USA, sample size of 64), CHS (southern Han Chinese individuals, China, sample size of 56), PEL (Peruvian individuals from Lima, Peru, sample size of 58) and YRI (Yoruba individuals from Ibadan, Nigeria, sample size of 38).

**Genetic map and no recombination regions.** We used the Decode genetic map in this study, which has been obtained by tracking more than 2,000 meioses in Islandic lineages Kong *et al*. (2002). The map was downloaded from the Table tool on the UCSC genome browser website Genome Bioinformatics Group of UC Santa Cruz (2013). We extract from the map regions having a recombination rate of exactly 0. There were 22,321 such regions, of varying lengths (see Supporting Information Figure S10), with the most common length being 10kb (6,457 regions) and mean length being around 48kb. We did not use HapMap recombination maps, as those are obtained using statistics of linkage disequilibrium (LD), which are affected by demography. In particular regions of high LD can be suggestive of either a low local recombination rate, or a short gene-genealogy of the sample used for LD computation, or both. So, by extracting regions of low “recombination rate” in LD-based genetic map, we might enrich the chosen regions in small underlying gene-genealogies, hence leading to the inference of a smaller population size. We saw this effect when applying Popsicle to regions extracted using HapMapCEU with a total recombination threshold of 10^−5^ Morgans per region (Supporting Information Figure S11).

**Comparison with PSMC.** Since its publication, the PSMC method has been widely used to estimate past population size over time in a number of organisms. Thus, it is important to assess how our reconstruction methods compares to the results of PSMC. We use the sequences of the parents, with missing genotypes imputed by BEAGLE, and cut the sequences into regions of 100 base pairs, as it is done in the original paper. If no pairwise difference is observed within a region between the pairs of alleles at the 100 base pairs, the region is considered homozygote. If at least one pairwise difference is observed, the region is considered heterozygote. PSMC is developed as a Hidden Markov Model, where the hidden states are the coalescent-times of each region, while the observed states are the heterozygosity of the regions. It models recombination in the transition probabilities from one region to its neighbor. Intuitively, if a locus has many heterozygote regions, its underlying coalescent-time is going to be inferred as large, whereas if a locus contains mostly homozygote regions, the coalescent-time is inferred as small. Chromosomes are given as independent sequences and only autosomes are used. We present the results of PSMC per population, as an average of the PSMC results over all parents within each population. For running PSMC, we use the same time intervals as the human study in the original PSMC paper.

**Application of Popsicle.** We apply Popsicle on the 22,321 low recombining regions for the 4 populations, under 2 different settings: in the first setting, we reconstruct an effective population size profile for every individual and average the results across all individuals from the same population (we refer to that setting as ‘Popsicle 1’); in the second setting, we use Popsicle on sub-samples of size 5 and compute the average of the obtained *N*(*t*) estimates within each population (we refer to that setting as ‘Popsicle 5’). We use the 2-step procedure described in the subsection “Inferring gene-genealogies from polymorphism data”. Because PSMC also infers the local gene-genealogies when performing its MCMC computations, we also extracted the local gene-genealogies from PSMC’s decoding (option -d of the program) and apply Popsicle 1 to them. The results seem highly unstable, questioning the reliability of the inferred local gene-genealogies from PSMC (see Supporting Information Figure S12).

**Data availability.** Simulated data can be regenerated using the commands given in the Supporting Information. Data from the 1000 Genomes Project is available on the ftp server ftp://ftp.1000genomes.ebi.ac.uk/vol1/ftp/

## Acknowledgments

We thank Martin Lascoux and Michael G. B. Blum for helpful comments on the manuscript.

## Supporting Information

### Derivation of the 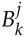

The relationship between the density function of the cumulative coalescent-times *π_k_* and the family of functions *q_j_* can be written under a matrix form. We define 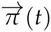 the vector of density functions of cumulative coalescent-times 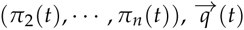 the vector (*q*_2_(*t*), ⋯, *q_n_*(*t*)) and the upper triangular matrix 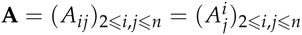. Then from equation 3, from Polanski *et al*. (2003) we have

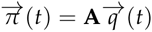

To prove that the 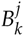 defined in theorem 1 can invert the relationship between *π_k_*(*t*) and *q_j_*(*t*), we show that the matrix **B** defined by 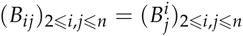 is the inverse matrix of **A**. We define 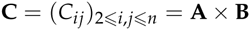. Our aim is to prove that **C** is in fact the identity matrix. First, we know that **C** is an upper triangular matrix, as both **A** and **B** are upper triangular matrices. To do so, we cover 4 separate cases: *C_in_* for 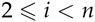 for 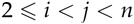, *C_ii_* for 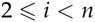 and finally *C_nn_*. For the computation of the two first cases, we need to introduce a notation:

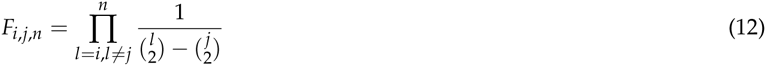

We know from partial fraction decomposition that

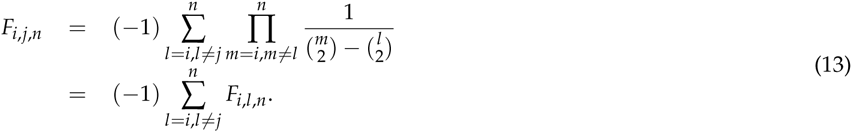

We compute the coefficients *C_in_*, for 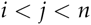:

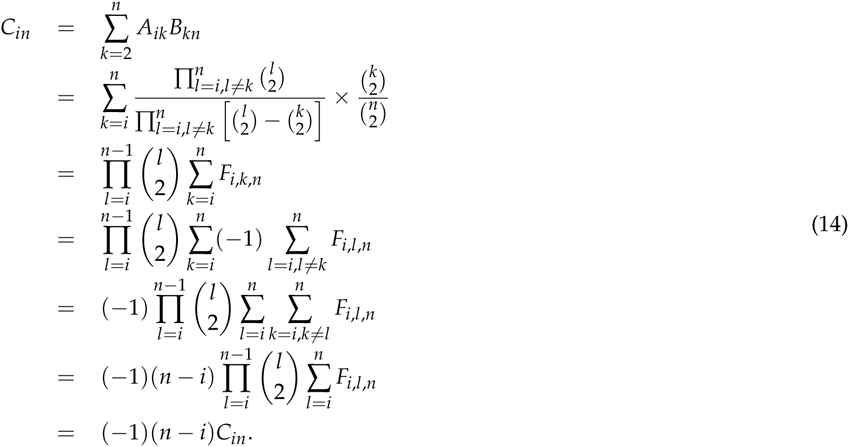

In the above calculation, we go from line 3 to line 4 by using equation 12. Then on the next line we exchange the two sums and by noticing that the terms under the *k*-indexed sum are not dependent on *k*, we obtain line 6. On line 6, we can notice that the factor after (−1)(*n* − *k*) is exactly the same as in line 3, thus is equal to *C_in_*. Since *n* ≠ *k*, only *C_in_* = 0 can satisfy *C_in_* =(*i* − *n*)*C_in_*.

We go on by computing our second case: the coefficients *C_ij_* for *i* < *j* < *n*:

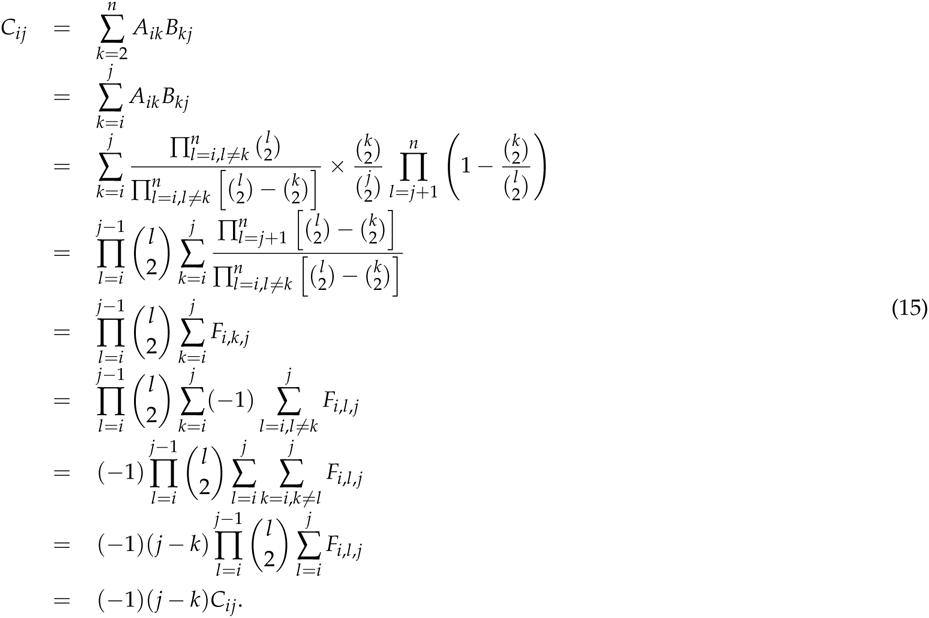

Similarly to the computation of *C_in_* above, the only way to satisfy *C_ij_* =(*i* − *j*)*C_ij_* for *i* < *j* < *n* is to have *C_ij_* = 0. Now, the remaining coefficients to be computed are the diagonal coefficients. For 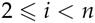:

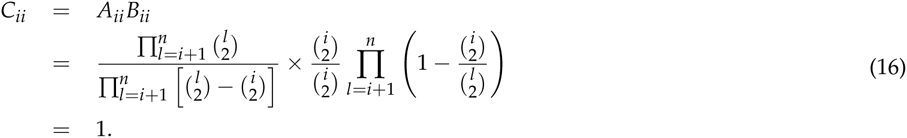

Finally,

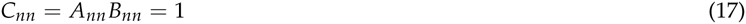

All the above computed coefficients prove that the matrix **C** is the identity matrix, hence **B** is the inverse matrix of **A**, which achieves to demonstrate theorem 1.

### The >ms commands for the simulations

All the times are given in units of 2 times the present haploid population size (see tab:s1 to tab:s4 for the exact values). The letter *n* can be replaced by any desired sample size.

- **scenario 1:** ms *n* 1 -t 1 -eN 0.025 2 -eN 0.25 0.5 -eN 0.5 1.5 -T
- **scenario 2:** ms *n* 1 -t 1 -G 6.93 -eG 0.2 0.0 -eN 0.3 0.5 -T
- **scenario 3:** ms *n* 1 -t 1 -G -0.732408192445406 -eG 1.5 0.0 -eN 2 4 -eN 3 3
- **scenario 4:** ms *n* 1 -t 1 -G 4605.17018598809 -eG 0.001 -2302.58509299405 -eG 0.002 0 -eN 0.003 0.2 -eN 0.0035 0.05 -eN 0.004 0.1

**Figure S1.**
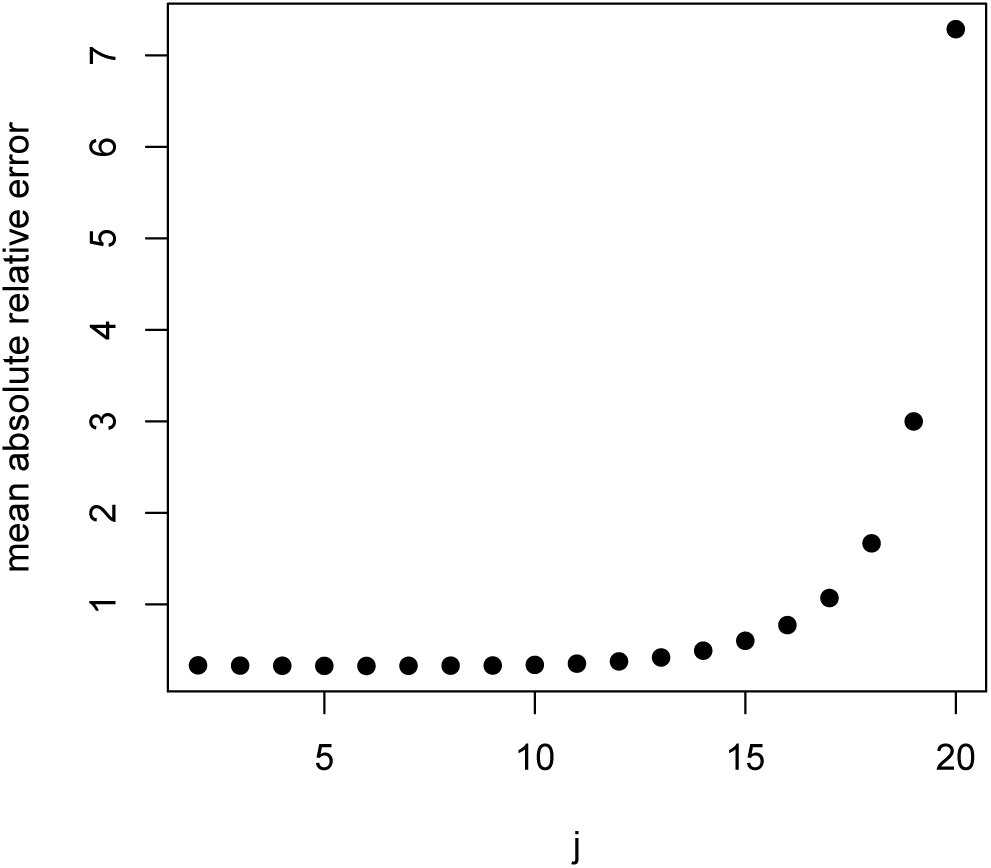
Accuracy of estimates of recent *N* as function of *j*. We compare estimates of *N* under scenario 1 with *n* = 20, between present and generation 1000 back in the past. Time is discretized in 100 equally sized bins and the accuracy of the *N* estimation is measured by the average relative error (see equation 10 in the main text).

**Figure S2.**
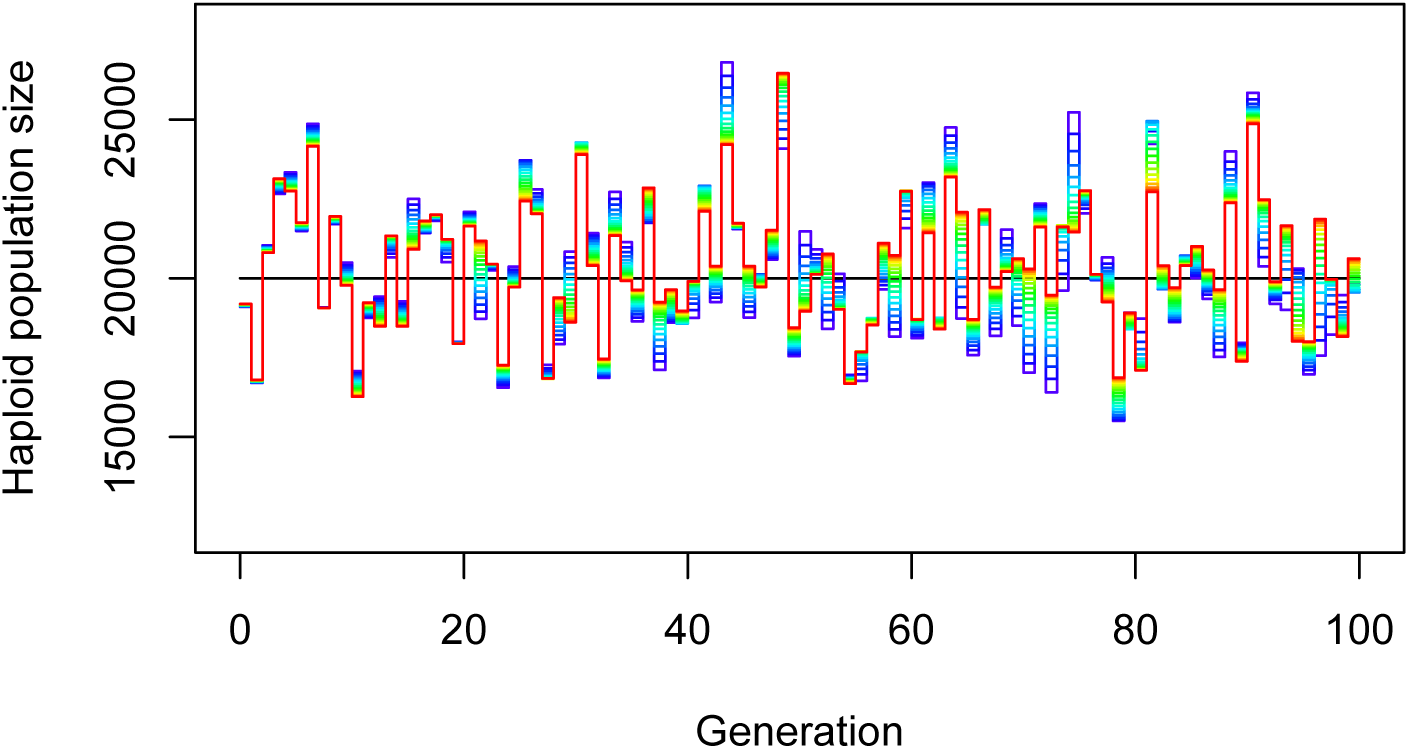
Estimation of *N*(*t*) depending on *j* during the first generations, scenario 1. Different values of *j* are indicated by the color of the solid lines, with a rainbow gradient from red (*j* = 2) to dark blue (*j* = 20).

**Figure S3.**
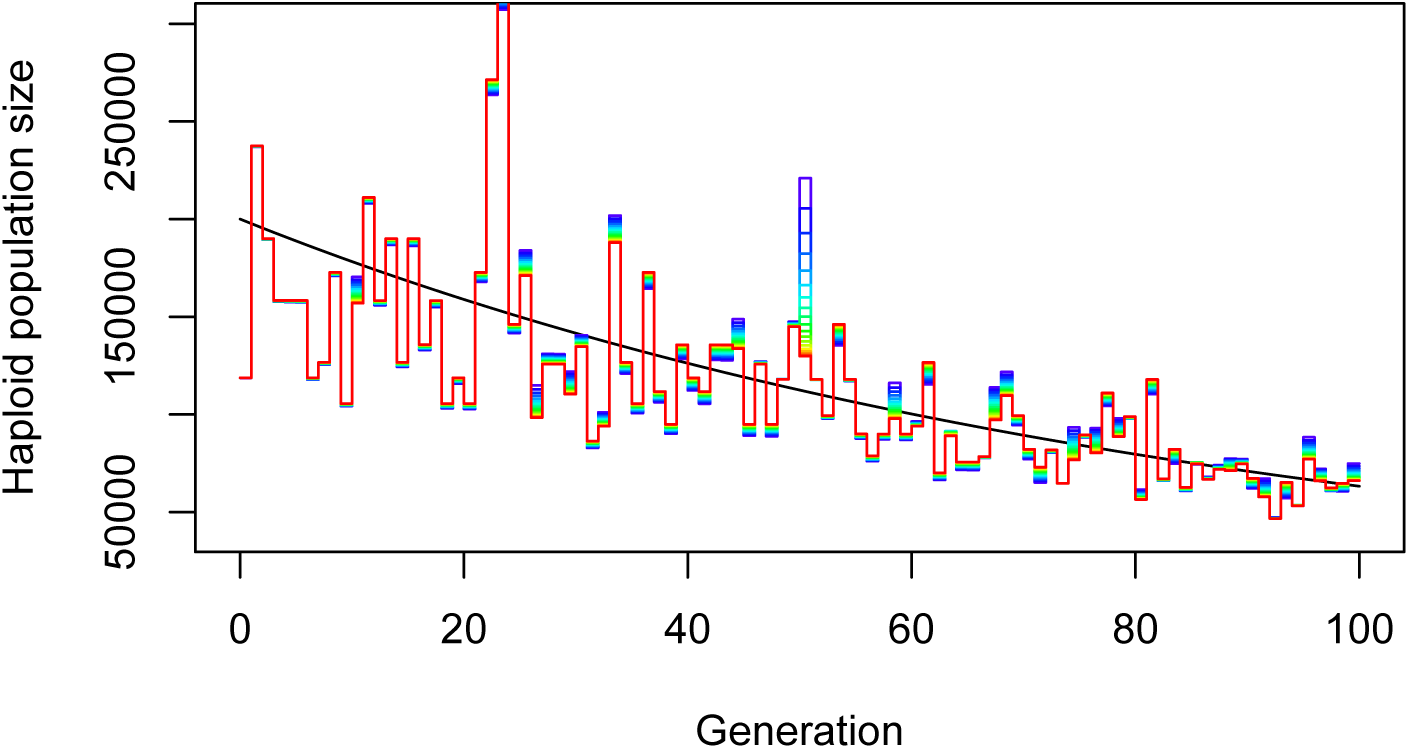
Estimation of *N*(*t*) depending on *j* during during the first generations, scenario 4. Different values of *j* are indicated by the color of the solid lines, with a rainbow gradient from red (*j* = 2) to dark blue (*j* = 20).

**Figure S4.**
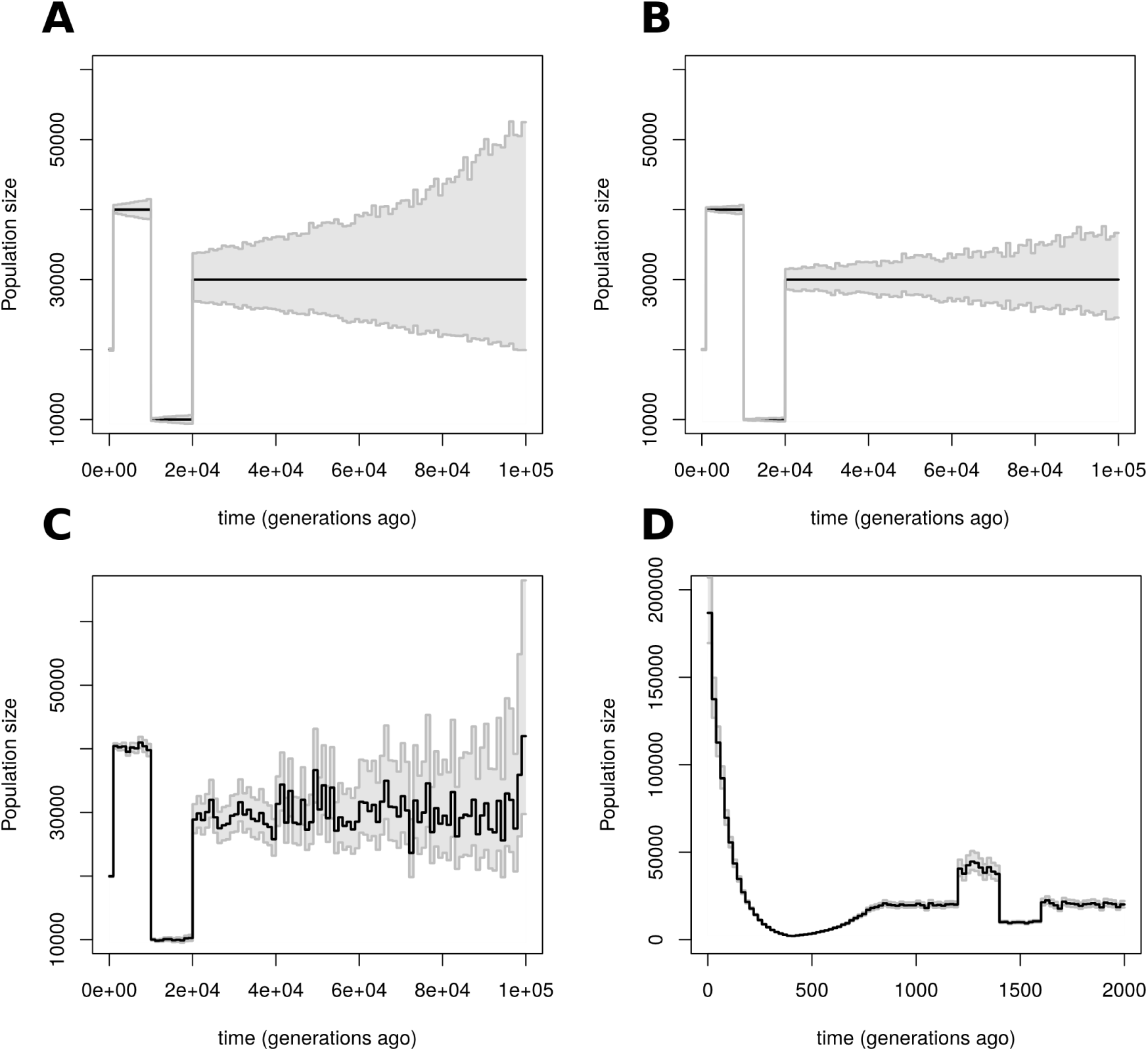
Uncertainty on the estimates of *N*(*t*). Results obtained by first simulating 1,000,000 independent gene-genealogies from model 1 with 20 haploid gene-copies and then (A) apply the theorem 10,000 times using 10,000 randomly sampled gene-genealogies from the 1,000,000 genealogies, or (B) apply the theorem 10,000 times using 50,000 randomly sampled gene-genealogies from the 1,000,000 genealogies. (C) Bootstrap results for model 1 using 20,000 gene-genealogies and 10,000 bootstrap replicates. (D) Bootstrap results for model 4 using 20,000 gene-genealogies and 10,000 bootstrap replicates. Time is discretized into 100 equally long intervals. We marked by a two solid gray lines the 2.5 and 97.5 percentiles of the 10,000 estimates of *N* within each interval. For (A) and (B), the black solid line represents the true value of *N*(*t*). For (C) and (D), the black solid line represents the reconstructed *N*(*t*) profile using our method on the 20,000 independent gene-genealogies.

**Figure S5.**
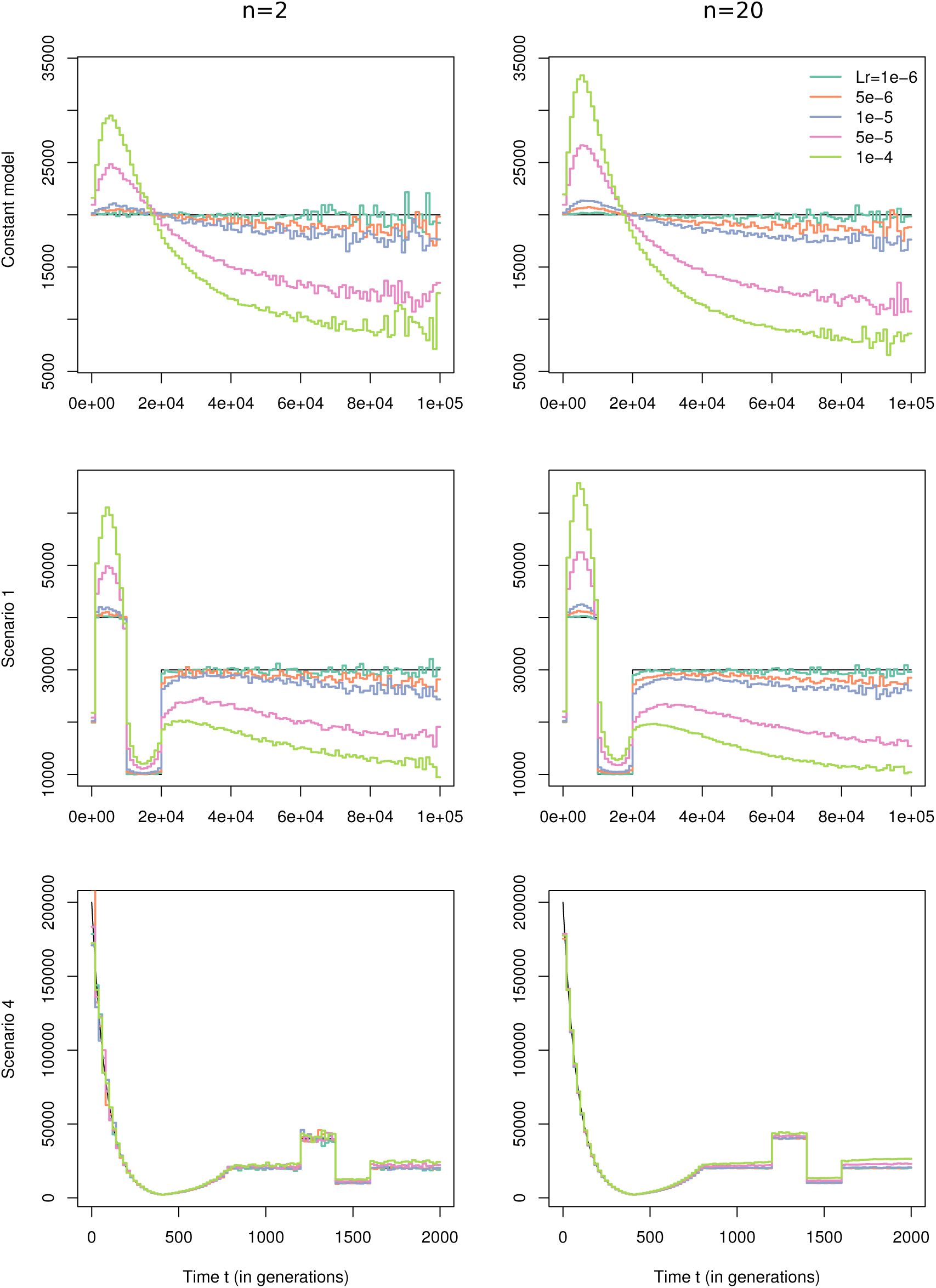
Effect of ignoring recombination. Comparison between *N*(*t*) reconstructed using average trees and true *N*(*t*) (black lines). The left column and right column display the results for samples of size 2 and 20 respectively, for different demographic scenarii: the constant size model (top line), scenario 1 (middle line) and scenario 4 (bottom line). The different cryptic recombination rates for each locus (in Morgan) is indicated by different colors and the values of the recombination of the segments are given in the legend.

**Figure S6.**
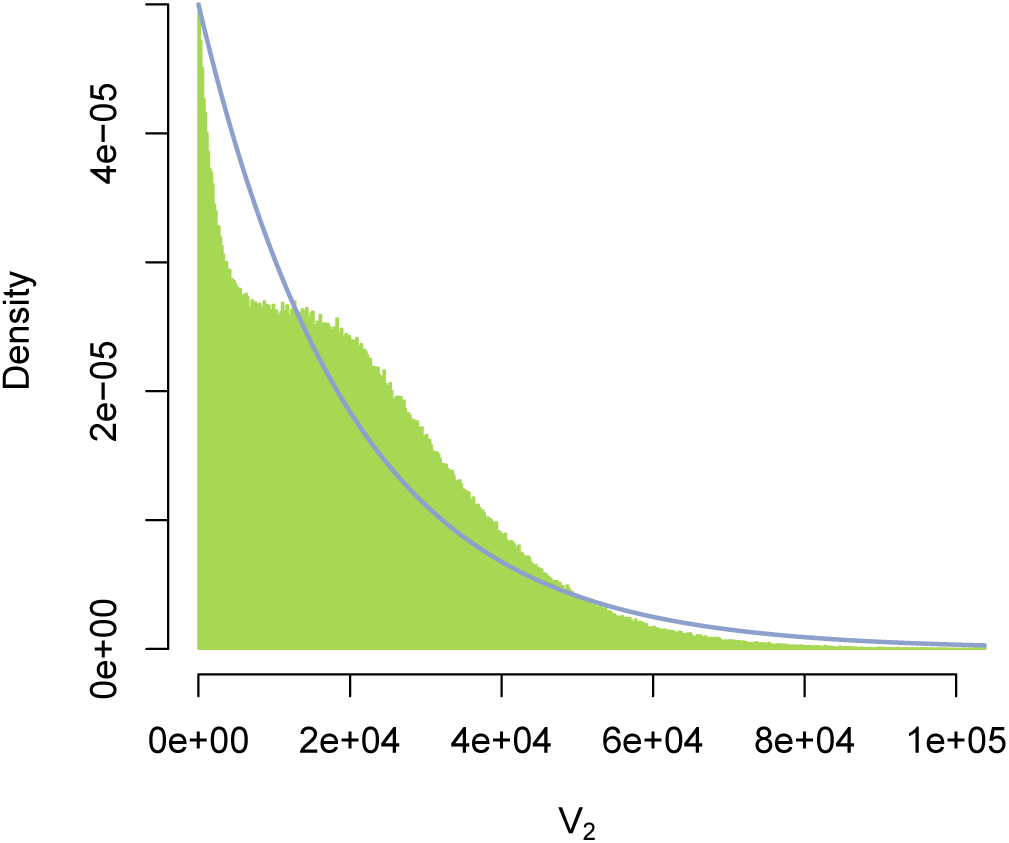
Density of *V*_2_ with cryptic recombination. Comparison between the expected density of *V*_2_ under the constant model for *n* = 2 (solid blue line) and the observed density of *V*_2_ under the constant model with recombination of *Lr* = 10^−4^ in green.

**Figure S7.**
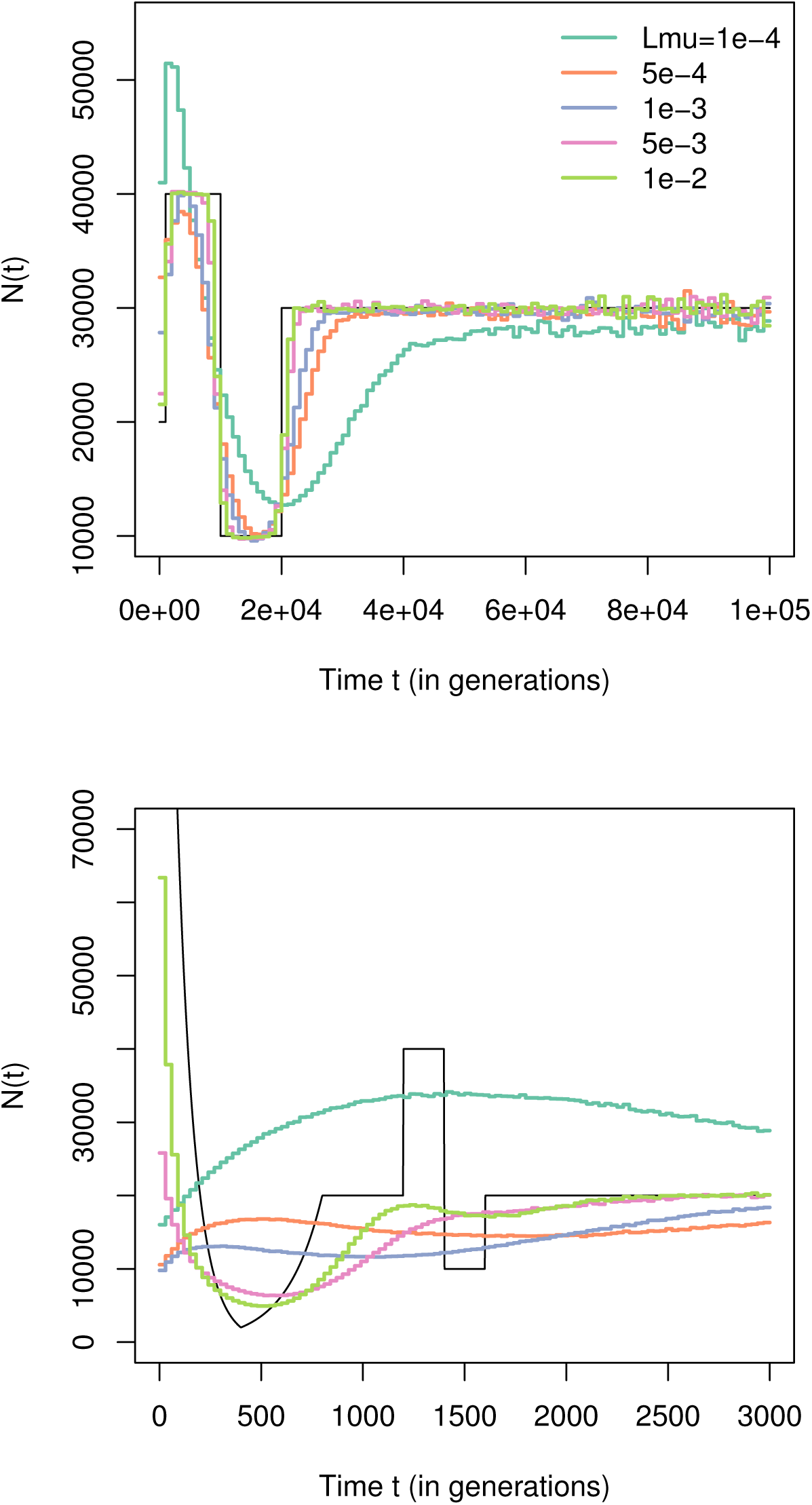
Effect of estimating trees from polymorphism data. Results of the 2 steps reconstruction method, applied with a sample size of 20, for 1,000,000 independent loci, evolving under scenario 1 (top figure) and scenario 4 (bottom figure). The mutation rate per locus *Lµ* is indicated by the color of the line and the legend gives the correspondence between the colors and the values.

**Figure S8.**
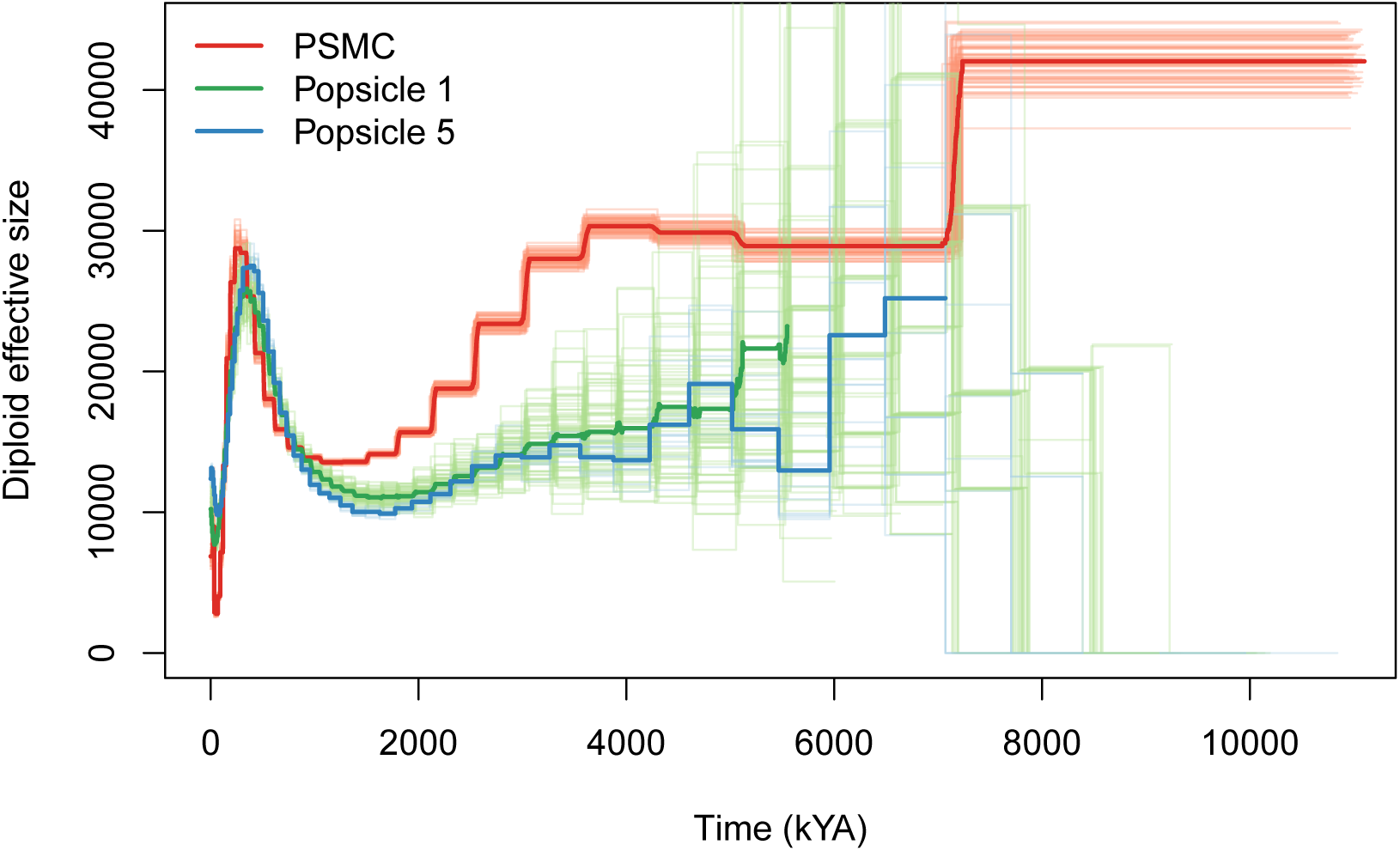
Comparison of methods on the CEU individuals. Zoomed out plot of the main figure 7.

**Figure S9.**
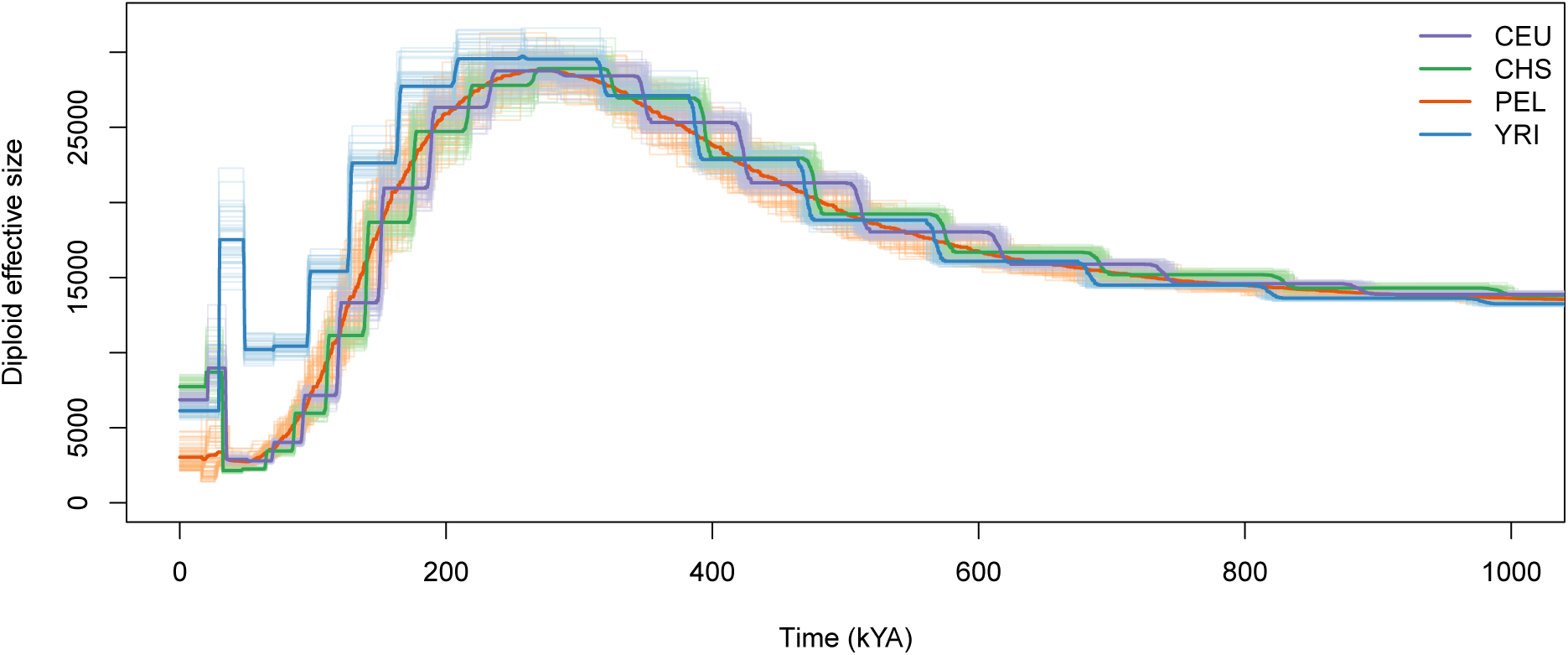
Results of PSMC on CEU, CHS, PEL and YRI. Thin light lines represent the population size reconstruction for one individual and thick lines indicate the average across individuals for a given population. Individuals from PEL have more variance in the estimated scaled mutation rate by PSMC, thus have time intervals that differ quite a bit from individual to individual when scaled back in years.

**Figure S10.**
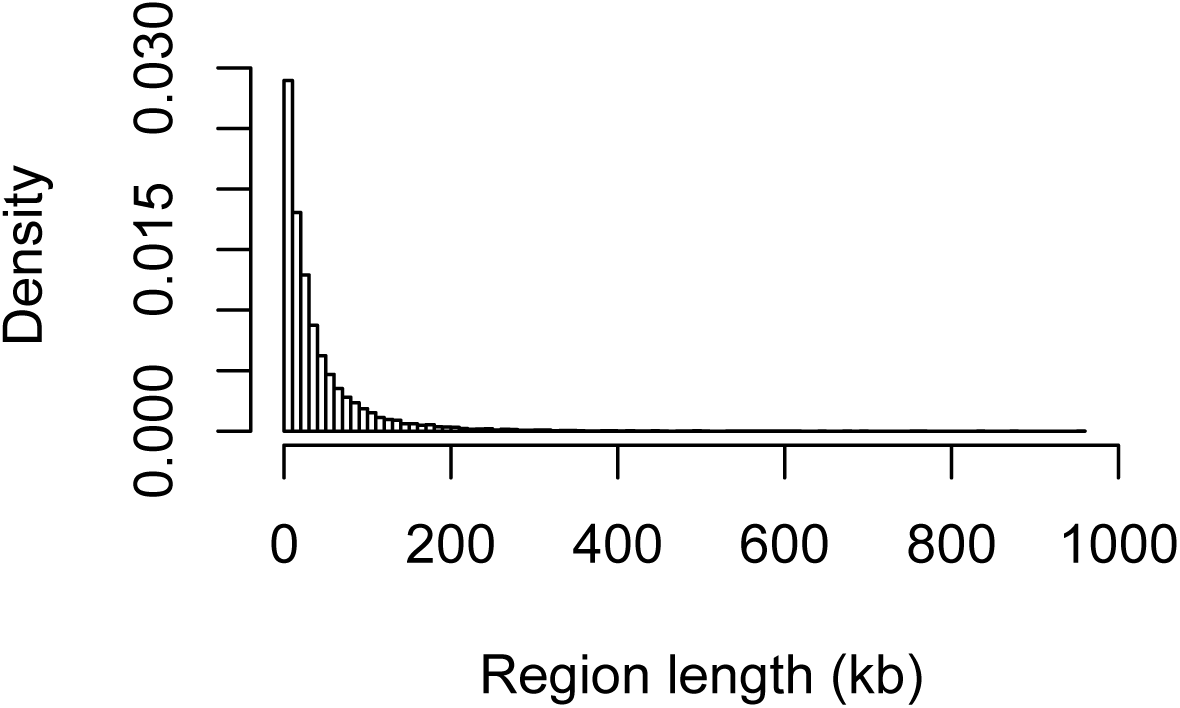
Distribution of length for the no recombining regions of the Decode genetic map.

**Figure S11.**
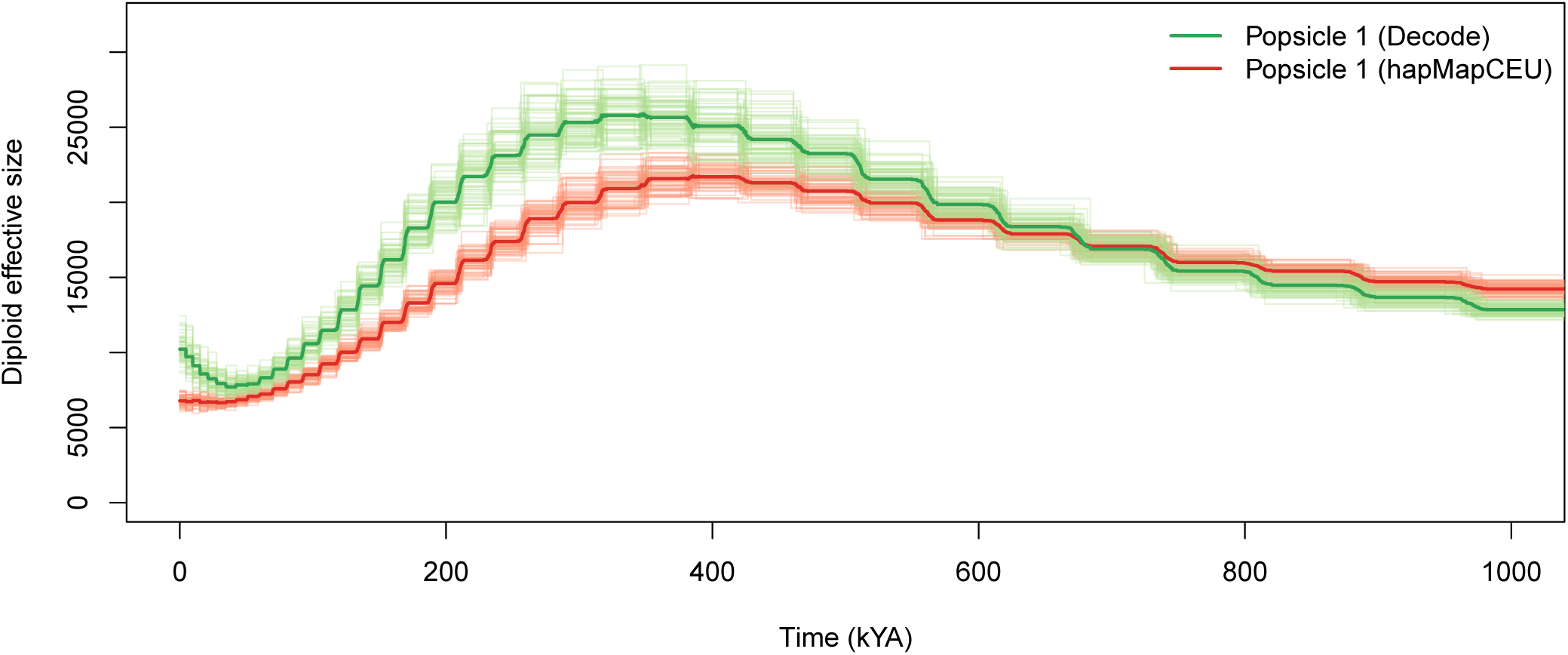
Comparison between Popsicle 1 using no recombining Decode regions (green lines) and Popsicle 1 using low recombining regions extracted from HapMapCEU. CEU samples.

**Figure S12.**
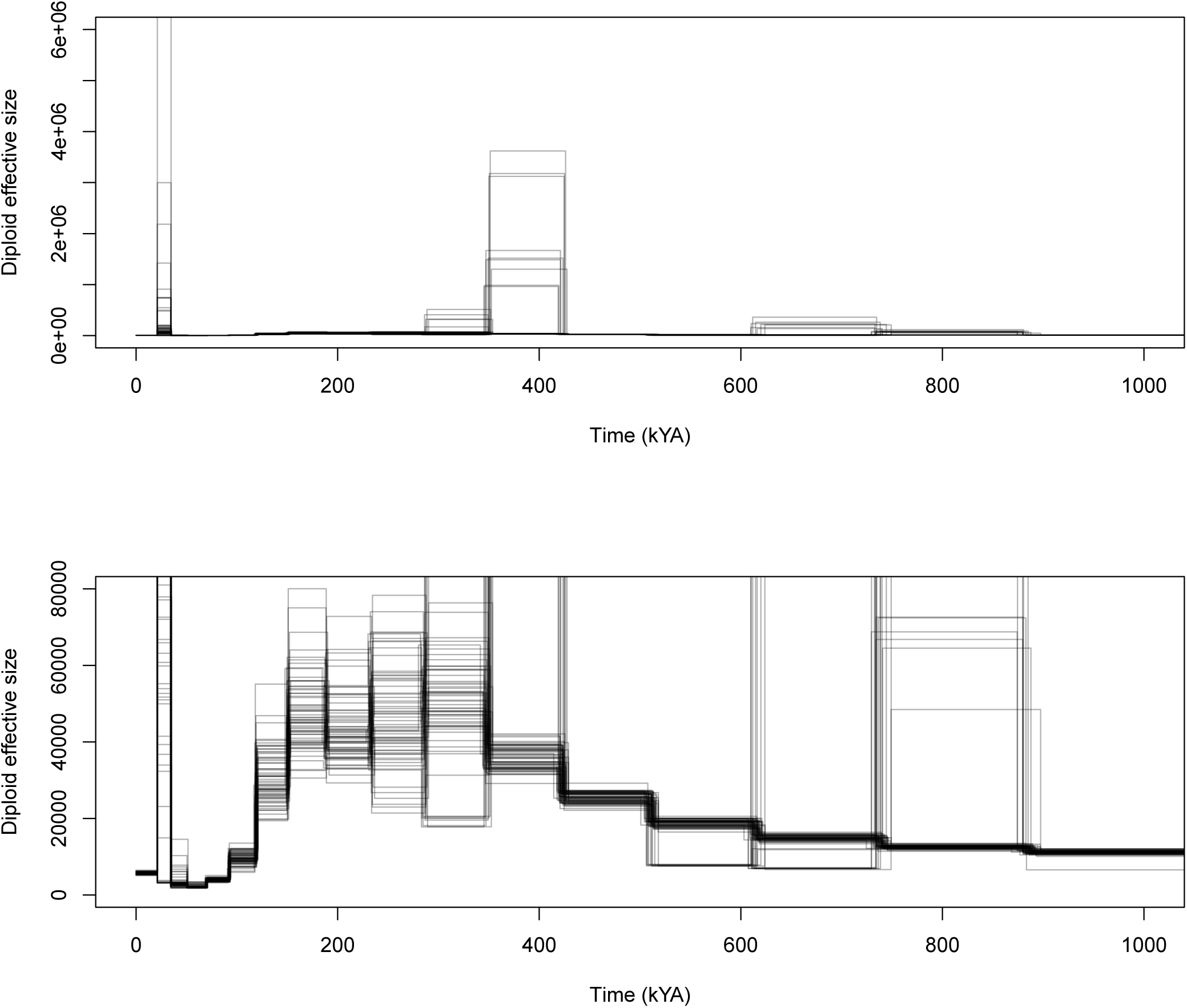
Application of Popsicle 1 to PSMC decoding gene-genealogies. Lower panel is a zoom in of the upper panel curve for smaller population size.

**Table S1.** Scenario 1

| Period (in gen.) | Haploid Size |
| --- | --- |
| 0-1,000 | 20,000 |
| 1,000-10,000 | 40,000 |
| 10,000-20,000 | 10,000 |
| > 20,000 | 30,000 |

**Table S2.** Scenario 2

| Period (in gen.) | Haploid Size | Parameters |
| --- | --- | --- |
| 0-16,000 | $N_0 \exp(-\alpha t)$ | $N_0 = 40,000, \alpha = 6.93/(2N_0)$ |
| 16,000-24,000 | 10,000 |  |
| > 24,000 | 20,000 |  |

**Table S3.**
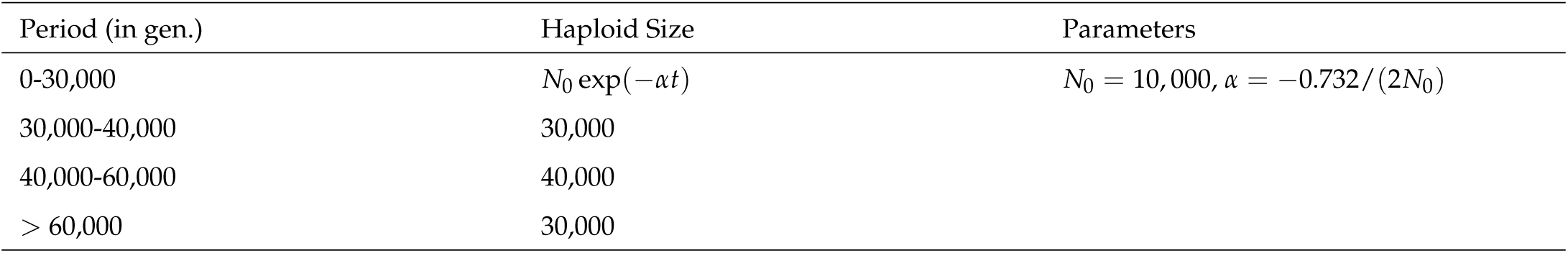
Scenario 3

| Period (in gen.) | Haploid Size | Parameters |
| --- | --- | --- |
| 0-30,000 | $N_0 \exp(-\alpha t)$ | $N_0 = 10,000, \alpha = -0.732/(2N_0)$ |
| 30,000-40,000 | 30,000 |  |
| 40,000-60,000 | 40,000 |  |
| > 60,000 | 30,000 |  |

**Table S4.** Scenario 4

| Period (in gen.) | Haploid Size | Parameters |
| --- | --- | --- |
| 0-400 | $N_0 \exp(-\alpha_1 t)$ | $N_0 = 200,000, \alpha_1 = 4605.2/(2N_0)$ |
| 400-800 | $N_1 \exp(-\alpha_2(t - 400))$ | $N_1 = 2,000, \alpha_2 = -2302.6/(2N_0)$ |
| 800-1,200 | 20,000 |  |
| 1,200-1,400 | 40,000 |  |
| 1,400-1,600 | 10,000 |  |
| > 1,600 | 20,000 |  |

